# BIN1 knockdown rescues systolic dysfunction in the aging heart

**DOI:** 10.1101/2022.08.23.504979

**Authors:** Maartje Westhoff, Silvia G. del Villar, Taylor L. Voelker, Phung N. Thai, Nipavan Chiamvimonvat, Eamonn J. Dickson, Rose E. Dixon

## Abstract

Cardiac dysfunction is a hallmark of aging in humans and mice. Here we report that a two-week treatment to restore youthful Bridging Integrator 1 (BIN1) levels in the hearts of 24-month-old mice rejuvenated cardiac function and substantially reversed the aging phenotype. Our data indicate that age-associated overexpression of BIN1 occurs alongside dysregulated endosomal recycling and disrupted trafficking of cardiac Ca_V_1.2 and type 2 ryanodine receptors. These deficiencies affect channel function at rest and their upregulation during acute stress. *In vivo* echocardiography revealed reduced systolic function in old mice. BIN1 knockdown using an adeno-associated virus serotype 9 packaged shRNA-mBIN1 restored the nanoscale distribution and clustering plasticity of ryanodine receptors and recovered Ca^2+^ transient amplitudes and cardiac systolic function toward youthful levels. Enhanced systolic function correlated with increased phosphorylation of the myofilament protein cardiac myosin binding protein-C. These results reveal BIN1 knockdown as a novel therapeutic strategy to rejuvenate the aging myocardium.

**Graphical Abstract:** 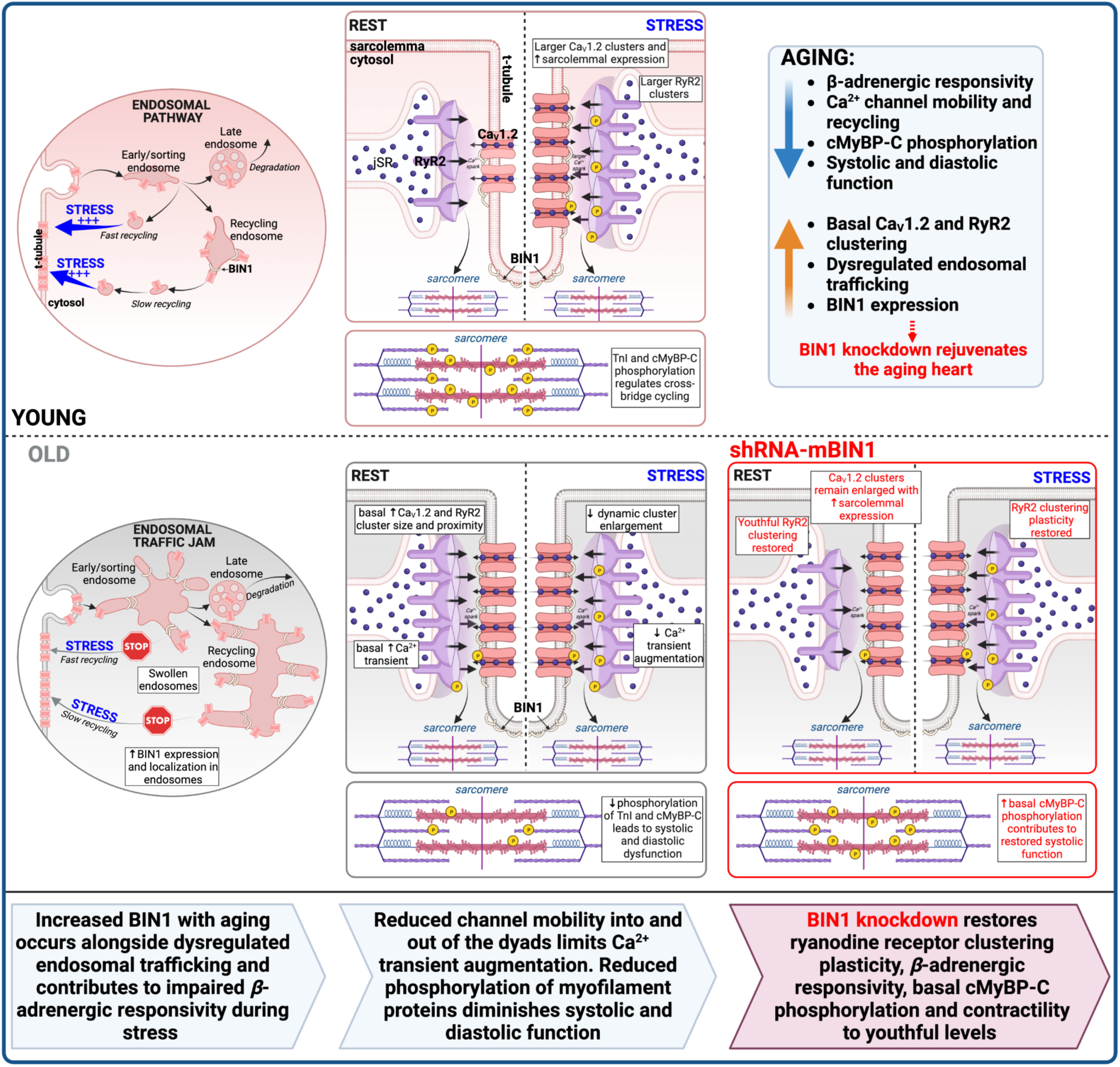

## Introduction

Old age is a major independent risk factor for cardiovascular disease which remains the leading cause of death globally^1, 2^. As humans advance into old age, regardless of overall health, certain intrinsic and progressive changes in cardiac structure and function occur including enhanced atrial and ventricular fibrosis, left ventricular hypertrophy, and diastolic dysfunction^3–5^. Maximal systolic function is also impaired due to a reduced responsiveness to sympathetic nervous system activation and subsequent *β-*adrenergic receptor (*β*-AR) signaling^3–5^. The decline of this reactive stimulus can precipitate exercise intolerance in the elderly and render the heart more vulnerable to damage during acutely stressful events when hypoxia and metabolic stress ensue^4, 6^. In mice, similar intrinsic cardiac changes occur over a much shorter lifespan and in the absence of other risk factors like smoking or diabetes, making mice a useful model to understand the molecular mechanisms of cardiac aging^3^.

The force of myocardial contraction and the rate and extent of the subsequent relaxation are tuned by both Ca^2+^ and myofilament-dependent processes. Examination and comparison of these points of regulation and their integration in the *in vivo* function of young and old mice may identify targets to improve the cardiac aging phenotype. Weakened myocardial contractile responses to *β*AR-stimulation have been linked to an age-associated decrease in the ability of the *β*AR signaling cascade to augment intracellular Ca^2+^ transient amplitude^7, 8^. This has been attributed to a reduced capacity to increase Ca_V_1.2 activity and availability, and thus to diminished Ca^2+^ influx during the action potential (AP)^5^. Activation of cAMP-dependent protein kinase A (PKA), downstream of *β*-AR stimulation, leads to phosphorylation of the channel complex^9, 10^, removing an inhibitory brake^11, 12^ and resulting in increased channel open probability (*P*_o_) due to potentiation of longer-duration ‘mode 2’ openings^13^, and an increased number of functional channels^14, 15^. We recently reported that *β*-AR activation also mobilizes a sub-sarcolemmal pool of Ca_V_1.2 channels, triggering a PKA-dependent, dynamic increase in sarcolemmal Ca_V_1.2 expression in ventricular myocytes^15, 16^. Consequent enlargement of Ca_V_1.2 channel clusters facilitates cooperative channel interactions^16, 17^, and contributes to an amplification of Ca^2+^ influx that rapidly tunes excitation-contraction (EC)-coupling to meet increased hemodynamic and metabolic demands. We hypothesized that loss of this rheostatic mechanism in aging myocytes could impose a narrow inotropic dynamic range on the system, limiting the response to *β*-AR stimulation.

Nanoscale redistribution and augmentation of cardiac Ca^2+^ channel clustering in response to *β*-AR stimulation also extends to RyR2. Accordingly, acute treatment with the *β*AR-agonist isoproterenol (ISO) or a phosphorylation-inducing cocktail promotes enhanced RyR2 cluster sizes^18, 19^, as recently reviewed^20^. This dynamic redistribution of RyR2 into larger clusters is orchestrated by bridging integrator 1 (BIN1)^18^ and is linked to increased Ca^2+^ spark frequency which summate to produce larger Ca^2+^ transients^19, 21^. There is widespread agreement that *β-*AR stimulated Ca^2+^ transient augmentation is impaired in the aging myocardium^5, 8^, however the underlying mechanisms are not understood.

Here, we present a comprehensive examination of cardiac Ca^2+^ channel function, their nanoscale distribution, and ISO-stimulated redistribution to test the central hypothesis that age-dependent disruption in the basal and on-demand trafficking and clustering of these channels restricts EC-coupling plasticity during myocardial aging and contributes to the pathophysiology of cardiac aging. We report that old myocytes exhibit basal super-clustering of both Ca_V_1.2 and RyR2, with no additional dynamic response to ISO. We link these deficits in cardiac Ca^2+^ channel recycling and mobility to an age-associated upregulation in BIN1 and the development of endosomal trafficking deficits or “endosomal traffic jams”. Crucially, we find that shRNA-mediated knockdown of BIN1 in old animals restores a young phenotype by re-establishing RyR2 organization, Ca^2+^ transient amplitude and their *β-*AR stimulated augmentation, and rejuvenating myofilament protein phosphorylation levels to recover youthful systolic function.

## Results

### β-AR stimulated I_Ca_ augmentation is diminished in aging

We began our study with an investigation into the Ca_V_1.2 channel response to *β-*AR stimulation by recording whole-cell Ca^2+^ currents (*I*_Ca_) from 3-month-old and 24-month-old (henceforth referred to as young and old, respectively) ventricular myocytes under control and 100 nM ISO-stimulated conditions. In young myocytes, ISO elicited a 1.59-fold increase in peak *I_Ca_* density (Fig. 1a-b, and Supplementary Table 1). However, old myocytes exhibited a blunted response showing only a 1.19-fold increase with ISO and basal currents were significantly larger in old myocytes than in young controls (Fig. 1a-c). The maximal ISO-stimulated current amplitude was similar in young and old myocytes (Fig. 1c), suggesting that the “ceiling” was intact but that the “floor” or baseline shifted with aging leaving less scope for current augmentation with acute adrenergic stress. Given the reduced ISO-stimulated current augmentation, these data suggest a reduced *β*-adrenergic responsiveness in old mice versus young.

**Figure 1.**
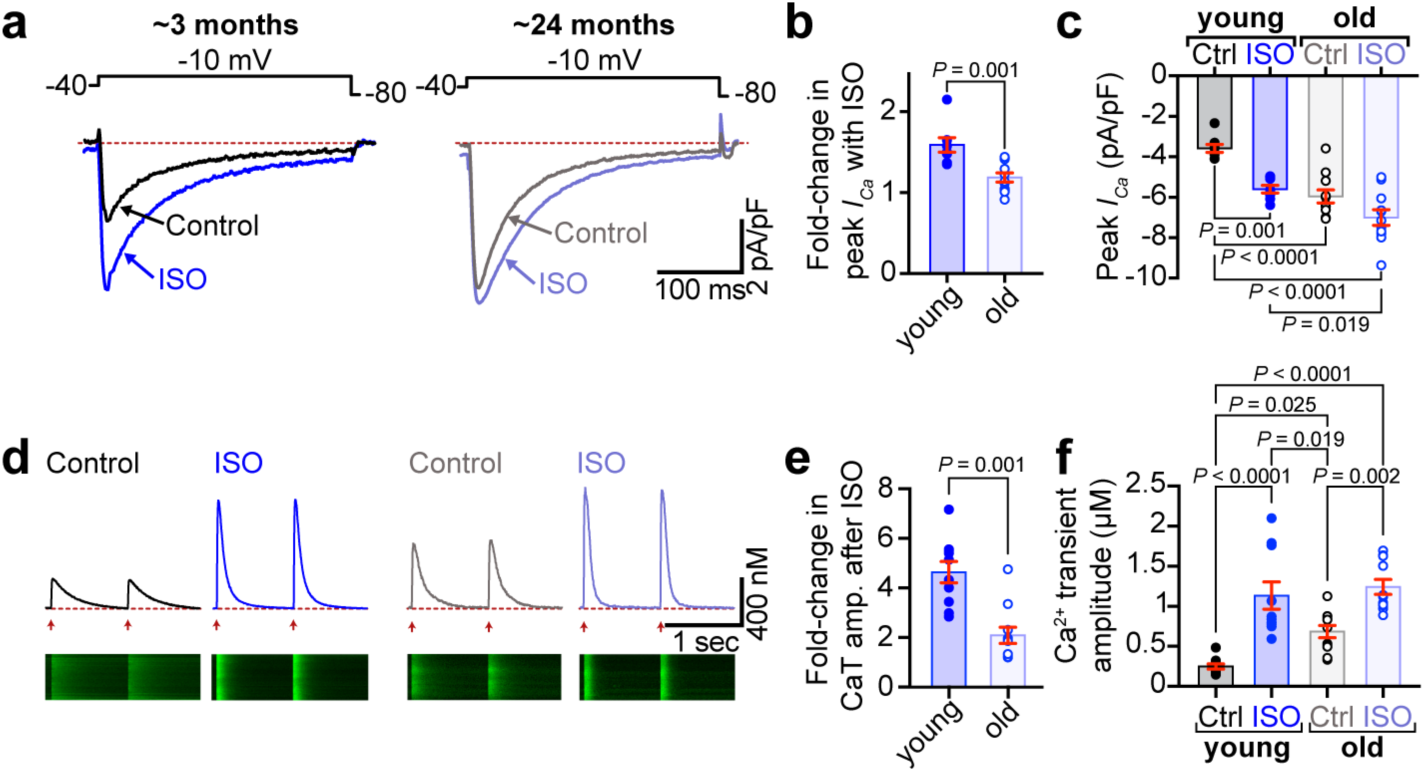
*β*-AR stimulated augmentation of *I*_Ca_ and Ca^2+^ transients is diminished in aging. **a**, representative whole-cell currents elicited from young and old ventricular myocytes before (control; black) and during application of ISO (blue). **b**, dot-plots showing the fold change in peak current with ISO in young (*N* = 6, *n* = 8) and old (*N* = 5, *n* = 11) myocytes. **c**, dot-plots showing peak *I_Ca_* before and after ISO. **d**, representative Ca^2+^ transients recorded before and after ISO from paced young (*N* = 6, *n* = 10) and old (*N* = 5, *n* = 11) myocytes. **e**, dot-plots showing the fold-increase in Ca^2+^ transient amplitude after ISO and **f**, Ca^2+^ transient amplitude before and after ISO. Unpaired Student’s t-tests were performed on data sets displayed in b and e. Two-way ANOVAs with post-hoc Tukey’s multiple comparison tests were performed on data displayed in c and f.

### Reduced capacity of β-AR stimulation to tune EC-coupling in aging myocytes

Influx of Ca^2+^ through Ca_V_1.2 channels during AP-mediated depolarization stimulates Ca^2+^-induced Ca^2+^-release from cross-dyad RyR2 to initiate EC-coupling. The amplitude of the resultant Ca^2+^ transient largely dictates the magnitude of myocardial contraction and the fraction of blood ejected, assuming constant Ca^2+^ sensitivity of the myofilaments. *β*-AR stimulation enhances both Ca_V_1.2 and RyR2-mediated Ca^2+^ influx to produce positive inotropy and a greater ejection fraction. We thus examined Ca^2+^ transients in young and old myocytes to investigate the effects of aging on *β-*AR tuning of EC-coupling. Transients evoked from Fluo-4AM-loaded myocytes paced at 1 Hz exhibited a 4.6 ± 0.4-fold enhancement in amplitude with ISO in young cells (Fig. 1d-f). In a parallel with the *I*_Ca_ results, basal Ca^2+^ transients in aging myocytes were larger than in young myocytes (Fig. 1d and f) and the fold-increase with ISO was almost halved (2.1 ± 0.3-fold change; Fig. 1e). These results further suggest reduced *β*-adrenergic responsiveness in old hearts and diminished myocardial responses to acute stress.

### Age-dependent alterations in nanoscale distribution and clustering of Ca_V_1.2 channels

Increased sarcolemmal Ca_V_1.2 channel expression and clustering at rest in aging cells could explain the age-dependent increase in basal *I*_Ca_. Furthermore, the reduced responsivity to ISO could be explained by an impaired capacity to mobilize/recycle additional endosomal Ca_V_1.2 channels in aging myocytes. In young cells, the result of ISO-stimulated channel recycling can be observed as Ca_V_1.2 super-clustering in single molecule localization microscopy (SMLM)^15^. To determine whether ISO-stimulated recycling was still functional in aging myocytes, we examined the nanoscale distribution and clustering of Ca_V_1.2 in t-tubule and crest regions of young and old myocytes using super-resolution SMLM with and without ISO-stimulation. These experiments confirmed that the super-clustering response to ISO was intact in the t-tubules of young myocytes, where Ca_V_1.2 channel cluster areas in ISO-treated young myocytes were 28.8% larger than unstimulated controls. However, in old myocytes, channels were already basally super-clustered, showed no augmentation in area with ISO, and were similarly sized to clusters in ISO-stimulated young cells (Fig. 2a-b). This phenomenon was isolated to the t-tubules as crest Ca_V_1.2 populations showed no significant alteration in cluster area (Supplementary Fig. 1). Overall, the age-dependent increase in basal Ca_V_1.2 channel cluster area and failure to elicit any additional increase with ISO suggests that a larger number of channels are already present in old myocyte t-tubule sarcolemma and that the ISO-stimulated channel insertion response is impaired in aging.

**Figure 2.**
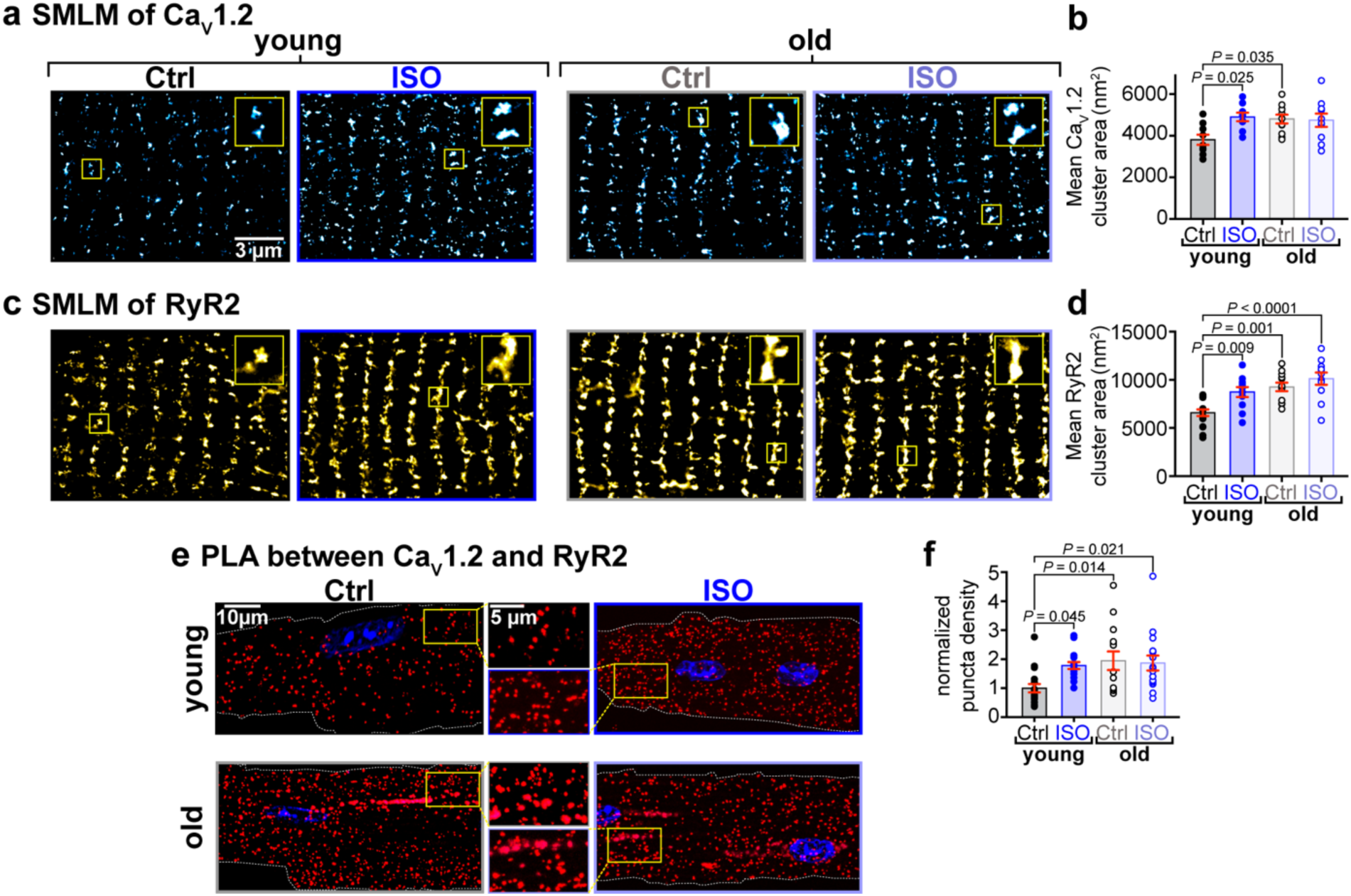
Basal super-clustering, impaired β-AR responsiveness, and enhanced proximity of Ca_V_1.2 and RyR2 in aged myocytes. **a**, SMLM localization maps showing Ca_V_1.2 channel localization and distribution in the t-tubules of young and old ventricular myocytes with or without ISO-stimulation. Yellow boxes indicate the location of the regions of interest magnified in the top right of each image. **b**, dot-plots showing mean Ca_V_1.2 channel cluster areas in young (control: *N* = 3, *n* = 9; ISO: *N* = 3, *n* = 9) and old (control: *N* = 4, *n* = 11; ISO: *N* = 4, *n* = 10) myocytes. **c** and **d**, show the same for RyR2 in young (control: *N* = 3, *n* = 17; ISO: *N* = 3, *n* = 11) and old (control: *N* = 3, *n* = 12; ISO: *N* = 3, *n* = 11) myocytes. **e**, Representative fluorescence PLA (red)/DAPI (blue) images of myocytes with and without ISO-stimulation. **f**, dot-plot summarizing the cellular density of PLA fluorescent puncta normalized to the density in young control cells (young control: *N* = 3, *n* = 19; young ISO: *N* = 3, *n* = 16; old control: *N* = 3, *n* = 14; old ISO: *N* = 3, *n* = 16). Statistical analyses on data summarized in b, d, and f were performed using two-way ANOVAs with post-hoc Tukey’s multiple comparison tests.

### Age-dependent alterations in nanoscale distribution and clustering of RyR2 channels

Other groups have reported a similar phosphorylation-induced cluster size expansion of the cross-dyad RyR2s in young ventricular myocytes^18, 19^. In agreement with those reports, SMLM performed on young myocytes immunostained against RyR2, revealed that average RyR2 cluster size grew by 32.7% in young myocytes after acute ISO treatment (Fig. 2c-d). However, mean RyR2 cluster areas in unstimulated old myocytes were similarly sized to the expanded clusters observed in ISO-stimulated young myocytes and did not undergo any further expansion with ISO. These findings suggest that the ISO-stimulated nanoscale redistribution of RyR2 is also impaired in aging.

### ISO-stimulated enhancement of Ca_V_1.2-RyR2 interactions is absent in aging

Since functional crosstalk between Ca_V_1.2 and RyR2 is an essential requirement for cardiac EC-coupling, we next examined the number of sites of proximity between Ca_V_1.2 and RyR2 using a proximity ligation assay (PLA). In this assay, a fluorescent signal is emitted at each location where Ca_V_1.2 and RyR2 come within 40 nm of each other^22^. In young myocytes, acute ISO treatment triggered the emergence of more Ca_V_1.2-RyR2 proximity sites (Fig. 2e-f). This phenomenon was not observed in old myocytes, where the number of Ca_V_1.2-RyR2 proximity sites appeared to have already reached a maximal level in the absence of any exogenous stimulation.

### Dynamic TIRF imaging reveals reduced ISO-stimulated Ca_V_1.2 insertion in aging

The basal super-clustering and absence of an additional response to ISO in old myocytes may indicate: (i) the endosomal reservoir is empty after already being recycled and inserted; and/or (ii) the trafficking and mobilization of channels is somehow impaired by aging. To interrogate these hypotheses, we visualized real-time dynamics of the channels utilizing a Ca_V_1.2 “biosensor” approach pioneered by our lab. In this technique, adeno-associated virus serotype 9 (AAV9)-Ca_V_β_2_-paGFP auxiliary subunits are transduced via retro-orbital injections and then visualized in isolated myocytes using TIRF microscopy as previously described^15, 16^. The relative balance between insertion and endocytosis dictates the overall expression of any membrane protein. An insertion-heavy mismatch between them will lead to increased membrane expression of the protein, while an endocytosis-heavy mismatch will favor reduced expression over time. We used image math to calculate the relative population of channels that were inserted during the ISO treatment, removed/endocytosed, or that remained static during the whole recording. As expected based on our previous studies^15, 16^, young myocytes responded to ISO by inserting more Ca_V_1.2 channels into the sarcolemma than they endocytosed, resulting in a net change in Ca_V_β_2_-paGFP in the TIRF footprint that on average amounted to 18% (Fig. 3a, c-g). However, old myocytes exhibited a comparatively blunted response with fewer ISO-stimulated insertions (Fig. 3b-c) and a larger population of static channels in the TIRF footprint suggesting impaired mobility (Fig. 3e). Consequently, the ISO-induced change in TIRF footprint Ca_V_β_2_-paGFP amounted to only 9.5% (Fig. 3g). Overall, these data reveal an age-dependent deficit in ISO-triggered insertion of Ca_V_1.2, and a shift towards reduced channel mobility or “stranding” at the plasma membrane.

**Figure 3.**
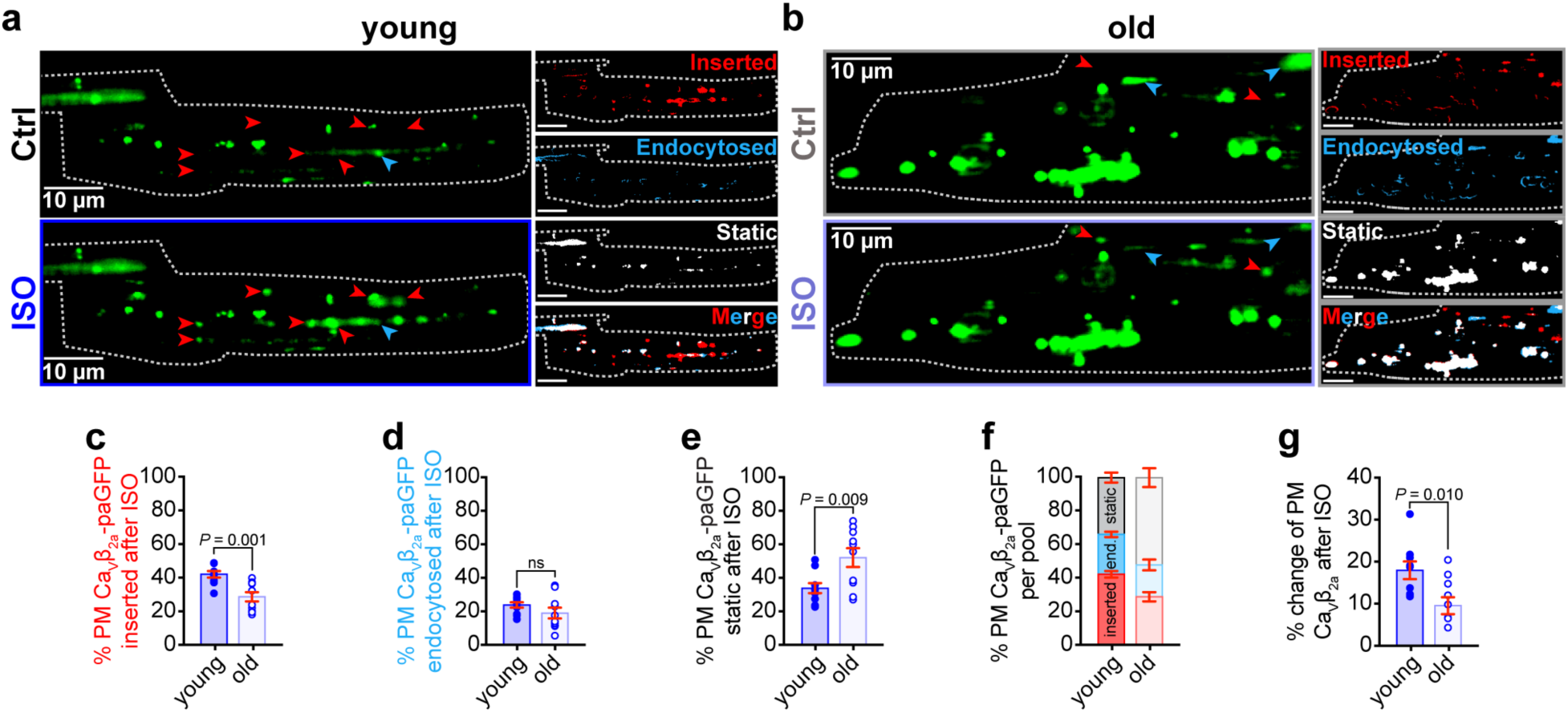
Dynamic TIRF imaging reveals age-associated trafficking deficits of Ca_V_1.2. **a** and **b**, Representative TIRF images of transduced Ca_v_β_2a_-paGFP young (**a**) and old (**b**) ventricular myocytes before (*top*) and after ISO (*bottom*). Channel populations that were inserted (red), endocytosed (blue), or static (white) during the ISO treatment are represented to the *right*. **c-e**, dot-plots summarizing the quantification of inserted (**c**), endocytosed (**d**), and static (**e**) Ca_v_β_2a_-paGFP populations. **f**, is a compilation of the data in c, d, and e to show the relative % of the channel pool that is inserted, endocytosed, and static. **g**, is a summary plot showing the net % change of plasma membrane Ca_v_β_2a_-paGFP after ISO for each cohort of young (*N* = 3, *n* = 10) and old (*N* = 3, *n* = 10) myocytes. Data were analyzed for statistical analysis using unpaired Student’s t-tests.

### Endosomal traffic jams impede mobilization of endosomal Ca_V_1.2 reservoirs in aging

We further investigated the effects of aging on the endosomal reservoir and capacity to mobilize Ca_V_1.2 channel cargo in response to ISO by examining EEA1-positive early endosome, and Rab11-positive recycling endosome pools of Ca_V_1.2 in young and old myocytes. Accordingly, Airyscan super-resolution imaging on young myocytes revealed immunostained Ca_V_1.2 on 14.89% of EEA1 positive pixels (Fig. 4a-b). As we have previously reported, *β*-AR activation stimulates the recycling of a portion of the early endosome Ca_V_1.2 to the plasma membrane via the Rab4-choreographed fast endosomal recycling pathway^15^. This was evidenced by a significant decrease in the % colocalization between EEA1 and Ca_V_1.2 after ISO in young myocytes. In old myocytes, there was a trending increase in the percentage of EEA1/Ca_V_1.2 colocalization in unstimulated cells and no ISO-induced endosomal emptying, suggesting trafficking alterations (Fig. 4b). Supporting this idea, EEA1-positive early endosomes were discernibly and quantifiably larger in old myocytes than in young (Fig. 4a and c).

**Figure 4.**
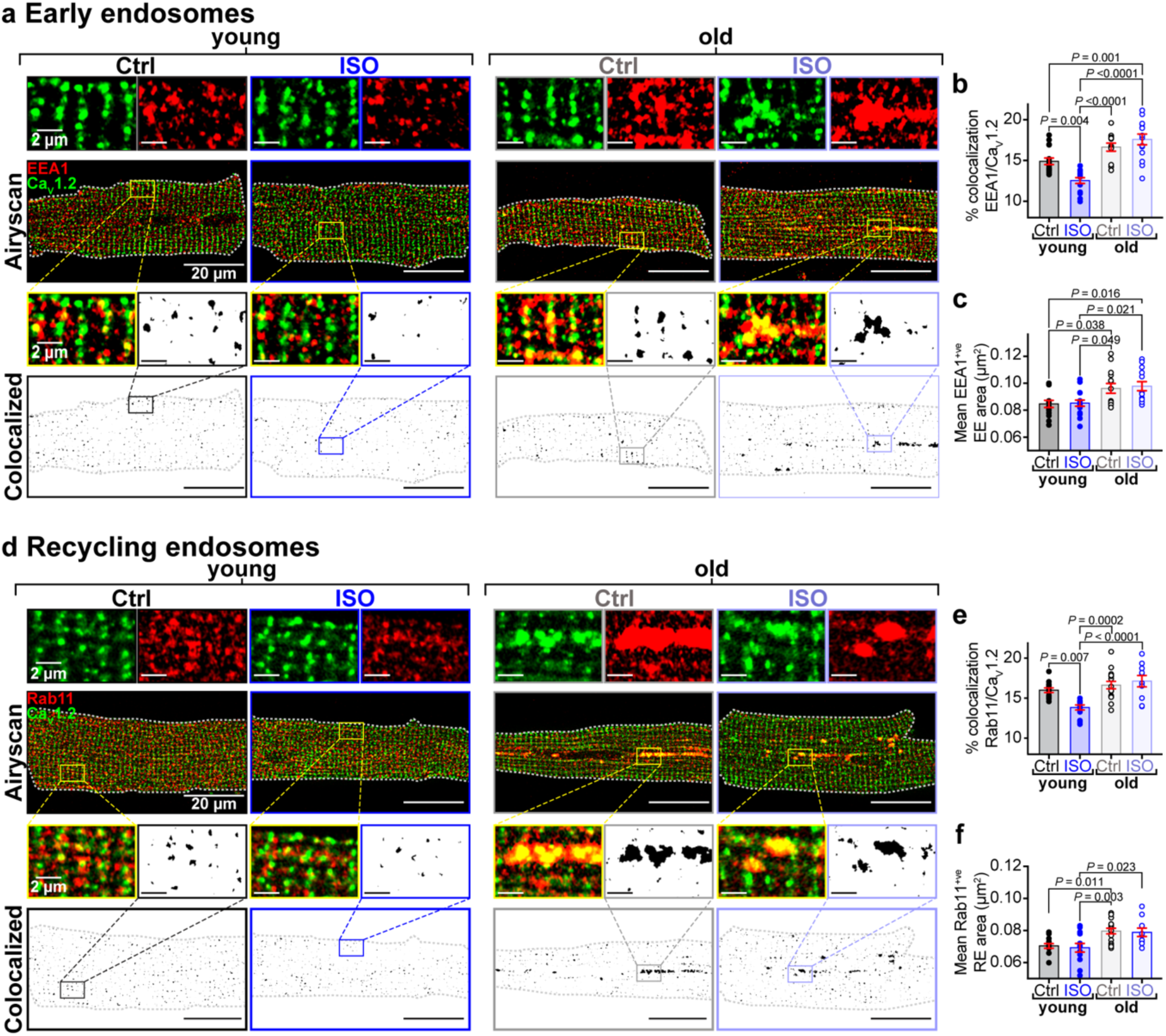
Aging impairs *β*-AR-stimulated Ca_v_1.2 recycling. **a**, Airyscan super-resolution images of Ca_V_1.2 and EEA1 immuno-stained myocytes with and without ISO. *Bottom:* Binary colocalization maps show pixels in which Ca_V_1.2 and EEA1 completely overlapped. **b**, dot-plots summarizing % colocalization between EEA1 and Ca_V_1.2, and **c**, EEA1 positive endosome areas in young (control: *N* = 3, *n* = 16; ISO: *N* = 3, *n* = 16) and old (control: *N* = 3, *n* = 14; ISO: *N* = 3, *n* = 14) myocytes. **d**-**f,** same layout for myocytes co-stained for Ca_V_1.2 and Rab11 recycling endosomes in young (control: *N* = 3, *n* = 13; ISO: *N* = 3, *n* = 13) and old (control: *N* = 4, *n* =16; old ISO: *N* = 3, *n* = 10) myocytes. Data were analyzed using two-way ANOVAs with post-hoc Tukey’s multiple comparison tests.

Recycling deficiencies extended into Rab11-positive recycling endosomes where a similar age-dependent absence of ISO-induced emptying of their Ca_V_1.2 cargo was observed in Rab11 and Ca_V_1.2 double-stained myocytes (Fig. 4d-e). Recycling endosomes were also found to be significantly enlarged in old myocytes compared to young (Fig. 4f). Rab7 positive late endosomes did not display any age-dependent alterations in their size or Ca_V_1.2 cargo suggesting that the degradation pathway may be intact (Supplementary Fig. 2).

### BIN1 is overexpressed and mislocalized in old myocytes

Endosomal enlargement has been linked to endosomal dysfunction and deficiencies in membrane protein recycling in Alzheimer’s disease (AD)^23^. Endosome swelling results from an imbalance in cargo coming into and leaving endosomes, sometimes referred to as endosomal traffic jams^24^. BIN1/amphiphysin II has been implicated as playing a role in the development of endosomal traffic jams in AD-stricken neurons^24^. In the heart, BIN1 is better known for its role in targeted delivery of Ca_V_1.2^25, 26^, t-tubule biogenesis^27^, micro-folding^28^, and maintenance^29^, and in dyad formation^30^. However, in neurons, where there are no t-tubules, BIN1 is known to play a role in mediating endosomal membrane curvature and tubule formation required for cargo exit and recycling from endosomes^31, 32^. BIN1 overexpression has been linked to aging and neurodegeneration in the brain^33–35^ and has been seen to produce endosomal expansion and traffic jams^24, 31, 32^. Furthermore, BIN1 expression levels are known to affect ion channel trafficking in both neurons^36^ and ventricular myocytes^25, 26^. We thus examined BIN1 expression and localization in young and old myocytes and found an age-associated upregulation (Fig. 5a-b) and redistribution of BIN1 with old myocytes exhibiting a vesicular, endosomal pattern of BIN1-staining alongside the expected t-tubule population (Fig. 5c; *left*).

**Figure 5.**
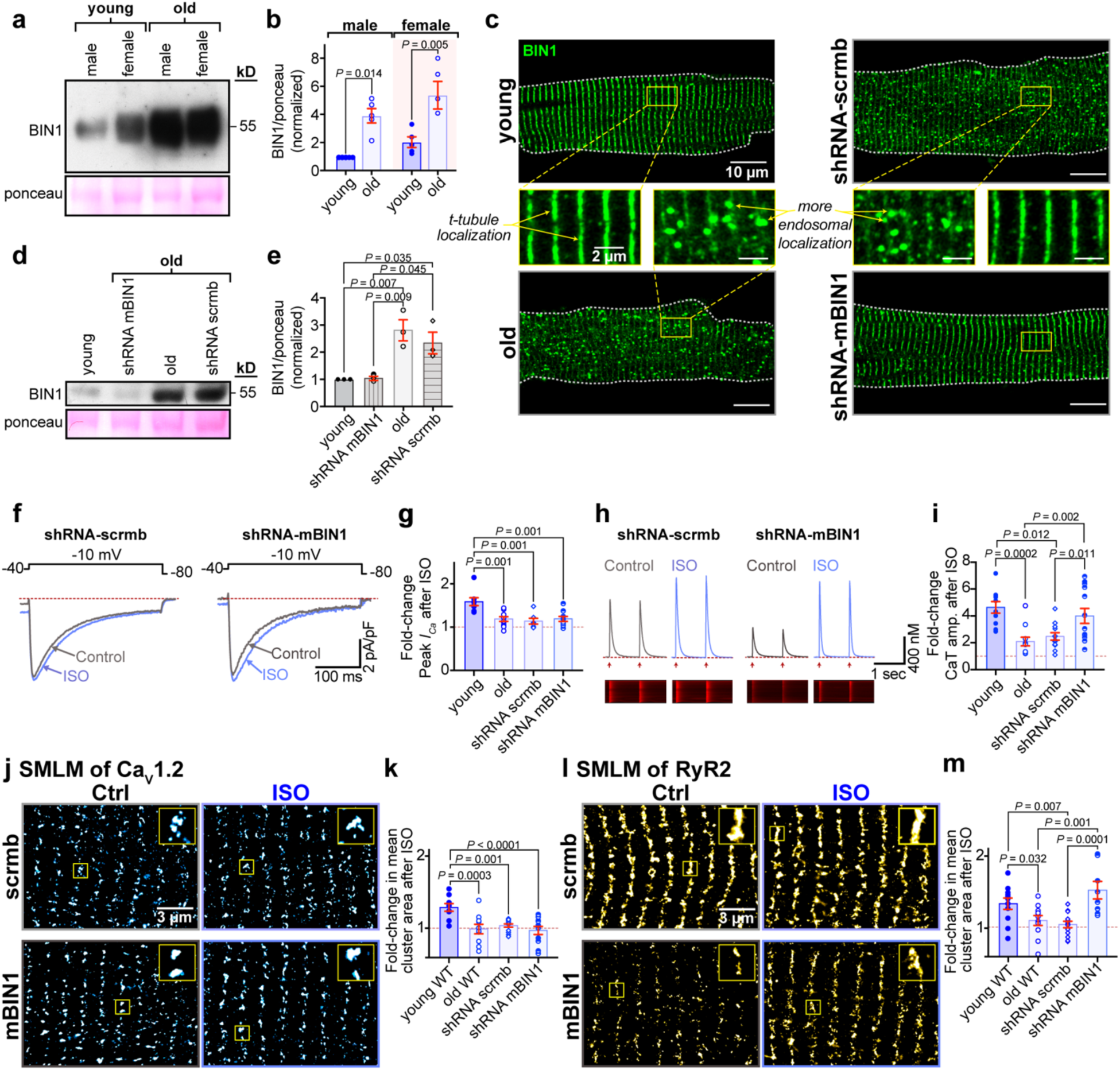
BIN1 knockdown restores *β*-AR augmentation of Ca^2+^ transients and RyR2 clustering dynamics. **a**, western blot of BIN1 expression in whole heart lysates from young and old mice. Ponceau was used for normalization. **b**, histogram showing normalized BIN1 levels relative to young male (*N* = 5, average of 1-2 replicates). **c**, representative Airyscan images of young, old, and old transduced myocytes immunostained against BIN1. **d**, western blot of BIN1 expression in whole heart lysates from: young, old, shRNA-mBIN1 or shRNA-scrmb transduced mice. Ponceau was used for normalization. **e**, histogram showing normalized BIN1 levels relative to young (*N* = 3, average of 1 - 3 replicates). **f**, representative whole-cell currents from shRNA-scrmb and shRNA-mBIN1 myocytes before and during ISO application. **g**, fold change in peak *I*_Ca_ with ISO in young, old, shRNA-scrmb (*N* = 3, *n* = 7) and shRNA-mBIN1 (*N* = 3, *n* = 9) myocytes. **h**, representative Ca^2+^ transients recorded from old shRNA-scrmb and shRNA-mBIN1 myocytes before and after ISO. **i**, fold increase after ISO from young, old, shRNA-scrmb (*N* = 3, *n* = 12) and shRNA-mBIN1 (*N* = 5, *n* = 13) myocytes. **j**, SMLM localization maps showing Ca_V_1.2 channel localization on t-tubules of myocytes from old shRNA-scrmb and shRNA-mBIN1, with or without ISO-stimulation. Regions of interest are highlighted by yellow boxes. **h**, fold change in mean Ca_V_1.2 channel cluster area with ISO in the old shRNA-scrmb (control: *N* = 3, *n* = 14; ISO: *N* = 3, *n* = 13) and shRNA-mBIN1 myocytes (control: *N* = 3, *n* = 12; ISO: *N* = 3, *n* = 11). **l** and **m**, show the same layout for RyR2 immunostained old shRNA-scrmb (control: *N* = 3, *n* = 12; ISO: *N* = 3, *n* = 13) and shRNA-mBIN1 myocytes (control: *N* = 3, *n* = 9; ISO: *N* = 3, *n* = 8). Old and young data points in g, i, k and m are reproduced from data in Figs. 1b, 1e, 2b and 2d respectively. Statistical analysis was performed on data in b using a two-way ANOVA, and data in e, g, i, k and m using one-way ANOVAs.

### BIN1 knockdown in old animals rejuvenates RyR2 plasticity and β-AR responsiveness

Since BIN1 became upregulated and mislocalized with aging with a more endosome-like pattern of expression, we hypothesized that knocking down BIN1 to young levels might resolve the endosomal traffic jams and restore *β*-AR responsivity to aging hearts. To test this, we transduced live mice with AAV9-GFP-mBIN1-shRNA (shRNA-mBIN1) or AAV9-GFP-scramble-shRNA (shRNA-scrmb; control) via retro-orbital injection. Two weeks post-injection, hearts were harvested, and successful shRNA-mediated knockdown of BIN1 was confirmed via both immunostaining and western blot analysis. BIN1 knockdown restored youthful localization of BIN1 to the t-tubules (Fig. 5c; *right*), and reduced BIN1 expression levels in old hearts to young heart levels (Fig. 5d-e). Control shRNA-scrmb transduced old hearts had similar BIN1 expression levels to non-transduced old hearts (Fig. 5d-e) and in all experiments, cells isolated from shRNA-scrmb transduced hearts maintained an old phenotype (Fig. 5c, 5f-m, Supplementary Fig. 3, and Supplementary Table 1). Contrary to our hypothesis, *I_Ca_* recordings from shRNA-mBIN1 transduced old myocytes did not indicate restoration of ISO responsivity (Fig. 5f-g, Supplementary Fig. 3a). In line with that finding, BIN1 knockdown did not reverse the endosomal swelling seen with aging suggesting that it is not an essential contributor to endosomal dysfunction in cardiomyocytes (Supplementary Fig. 4). However, basal Ca^2+^ transient amplitude in shRNA-mBIN1 transduced old myocytes resembled that of young myocytes and *β*-adrenergic responsiveness was fully restored to young levels (Fig. 5h-i, Supplementary Fig. 3b).

The recovery of Ca^2+^ transient amplitude suggested that BIN1 knockdown may predominantly impact RyR2 function and/or distribution and not Ca_V_1.2. To investigate that possibility, we performed SMLM on shRNA-mBIN1 transduced myocytes finding the nanoscale distribution of Ca_V_1.2 was unaltered compared to non-transduced old myocytes in that they remained basally super-clustered and did not recover the super-clustering response to ISO (Fig. 5j-k, and Supplementary Fig. 3c). In contrast, RyR2 appeared rejuvenated and exhibited a young cell-like nanoscale distribution and response to ISO (Fig. 5l-m, and Supplementary Fig. 3d). Overall, these results suggest that knockdown of BIN1 in aging hearts recovers cardiac *β*-adrenergic responsiveness by restoring RyR2 clustering plasticity.

### BIN1 knockdown in old animals rescues systolic but not diastolic function

Since all our experiments were performed in isolated cells, the question remained, would BIN1 knockdown improve old heart contractile function *in vivo*? To address this, we performed echocardiography in young, old, and old shRNA-transduced mice. In conscious mice fractional shortening (FS) and ejection fractions (EF) were reduced in old mice compared to young (Fig. 6a-c) suggesting an age-dependent systolic dysfunction. To examine the effects of shRNA-scrmb or shRNA-mBIN1 on old heart function, we performed echocardiograms on old mice to record their initial function prior to treatment with shRNA (Fig. 6d-e; *left*). Those mice were then retro-orbitally injected with either shRNA-scrmb (control) or shRNA-mBIN1 and after a two-week transduction period, repeat echocardiograms were obtained (Fig. 6d-e; *right*). In agreement with the rejuvenating effects of BIN1 knockdown observed in the *in vitro* functional analysis of Ca^2+^ transients, basal ventricular function was restored to a more youthful phenotype upon BIN1 knockdown, with shRNA-mBIN1 treated old mice displaying improved FS and EF (Fig. 6f-g).

**Figure 6.**
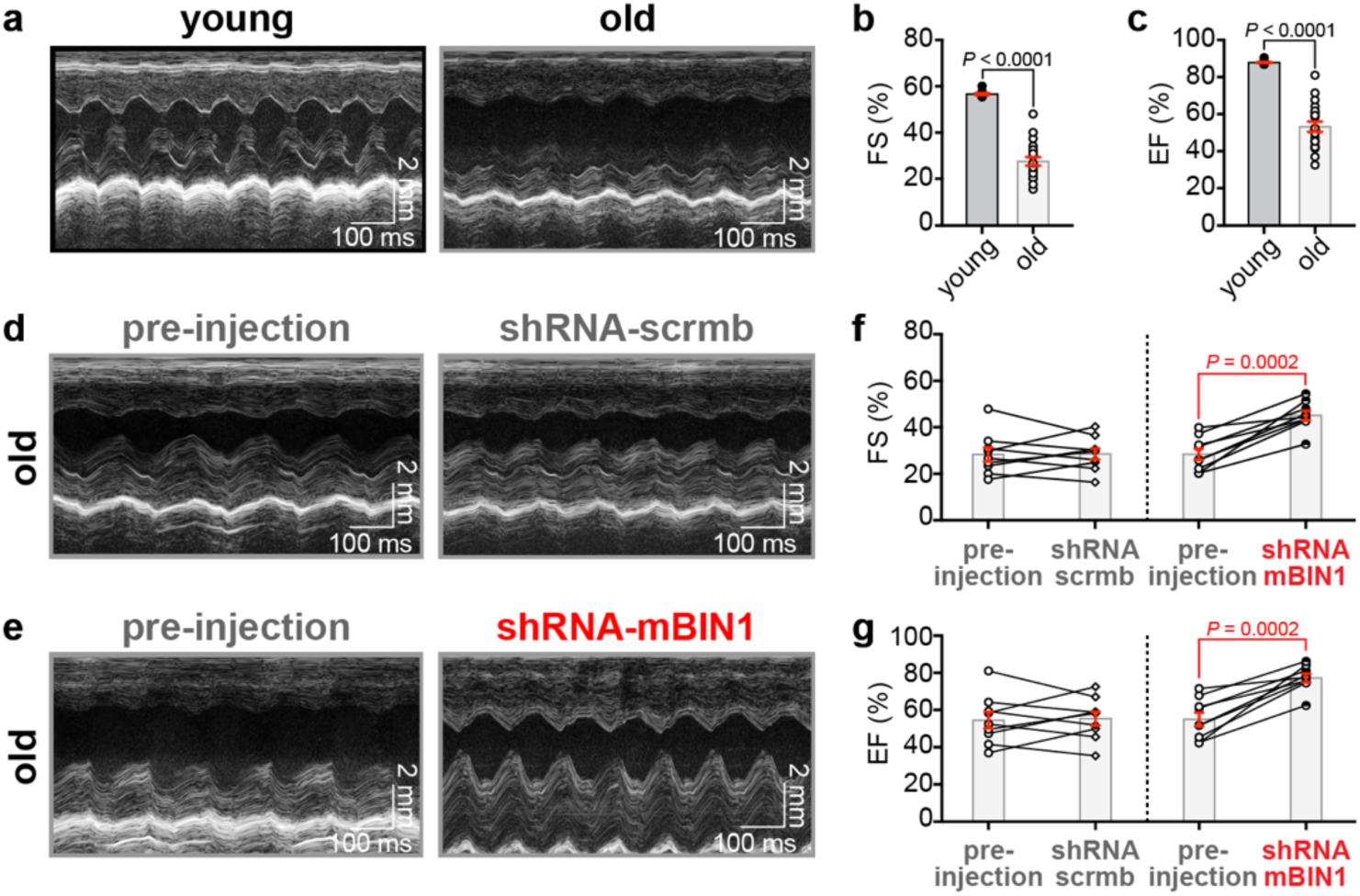
BIN1 knockdown improves cardiac contractility in old mice. **a**, representative M-mode echocardiogram images from conscious young and old mice. Summary dot-plots showing **b,** fractional shortening and **c,** ejection fraction in young (*N* = 11) and old (*N* = 20) mice are shown. **d** and **e,** representative M-mode echocardiogram images from conscious old mice before (*left*) and two-weeks after RO-injection (*right*) of shRNA-scrmb (**d**) and shRNA-mBIN1 (**e**). Summary dot-plots for **f,** fractional shortening and **g**, ejection fraction showing paired results before and after RO-injection for shRNA-scrmb (*N* = 9) and shRNA-mBIN1 (*N* = 9) groups are displayed. Unpaired Student’s t-tests were performed on data displayed in b and c. Paired Student’s t-tests were performed on data displayed in f and g.

Old mice also exhibited signs of left ventricular (LV) hypertrophy including increased LV mass (Supplementary Fig. 5a). Two weeks of BIN1 knockdown did not significantly improve LV mass although there was a downward trajectory toward a more youthful LV mass in paired old mice pre- and post-shRNA-mBIN1 that was not apparent in the shRNA-scrmb cohort (Supplementary Fig. 5b). LV end diastolic volumes (EDV) and end systolic volumes (ESV) were significantly increased in old mice compared to young (Supplementary Fig. 5e and g). The resultant increase in preload in old mice would be expected to enhance ventricular output via the Frank-Starling mechanism to maintain stroke volume in the face of reduced contractility. shRNA-mBIN1 transduction in old mice significantly reduced EDV and ESV to youthful levels (Supplementary Fig. 5f and h). Accordingly, cardiac output (CO) was significantly rejuvenated by BIN1 knockdown in old mice, increasing from a mean of 14.82 mL/min pre-injection to 17.32 mL/min two weeks post-knockdown and approaching the 18.61 mL/min seen in young mice (Supplementary Fig. 5i-j). The restoration of youthful CO, EDV, ESV, FS and EF by BIN1 knockdown suggests this treatment rejuvenates systolic function in old mice.

We further measured the *in vivo* effects of ISO on ventricular contractile function by performing echocardiograms on anesthetized young and old mice before and after intraperitoneal injection of ISO. As above, repeat echocardiograms were performed on old mice 2-weeks after retro-orbital injection of shRNA-scrmb or shRNA-mBIN1 (Supplementary Figs. 6 and 7). As has been previously reported, resting and ISO-stimulated heart rates were significantly lower in old mice compared to young^37^, but neither the scrambled shRNA nor the BIN1 knockdown had any significant effect on heart rate suggesting that pacemaker activity is not affected by these treatments (Supplementary Figs. 5c-d, 6a-b, and 7a-b). In agreement with the *in vivo* data from conscious mice, basal ventricular contractile function was rejuvenated by BIN1 knockdown with CO, FS, EF, EDV and ESV (Supplementary Figs. 6 and 7) all improving toward youthful levels. Importantly this effect was not observed in shRNA-scrmb transduced mice suggesting a specific effect of BIN1 knockdown.

To assess changes in diastolic function with aging, we performed pulsed wave Doppler imaging on unconscious mice. Old mice displayed an enhanced isovolumetric relaxation time (IVRT) and reduced mitral valve early to late ventricular filling velocity ratio (MV E/A), indicative of diastolic dysfunction, which was not remedied with BIN1 knockdown (Fig. 7). Altogether, these results suggest that BIN1 knockdown may represent a new therapeutic strategy to improve age-related deficits in systolic function.

**Figure 7.**
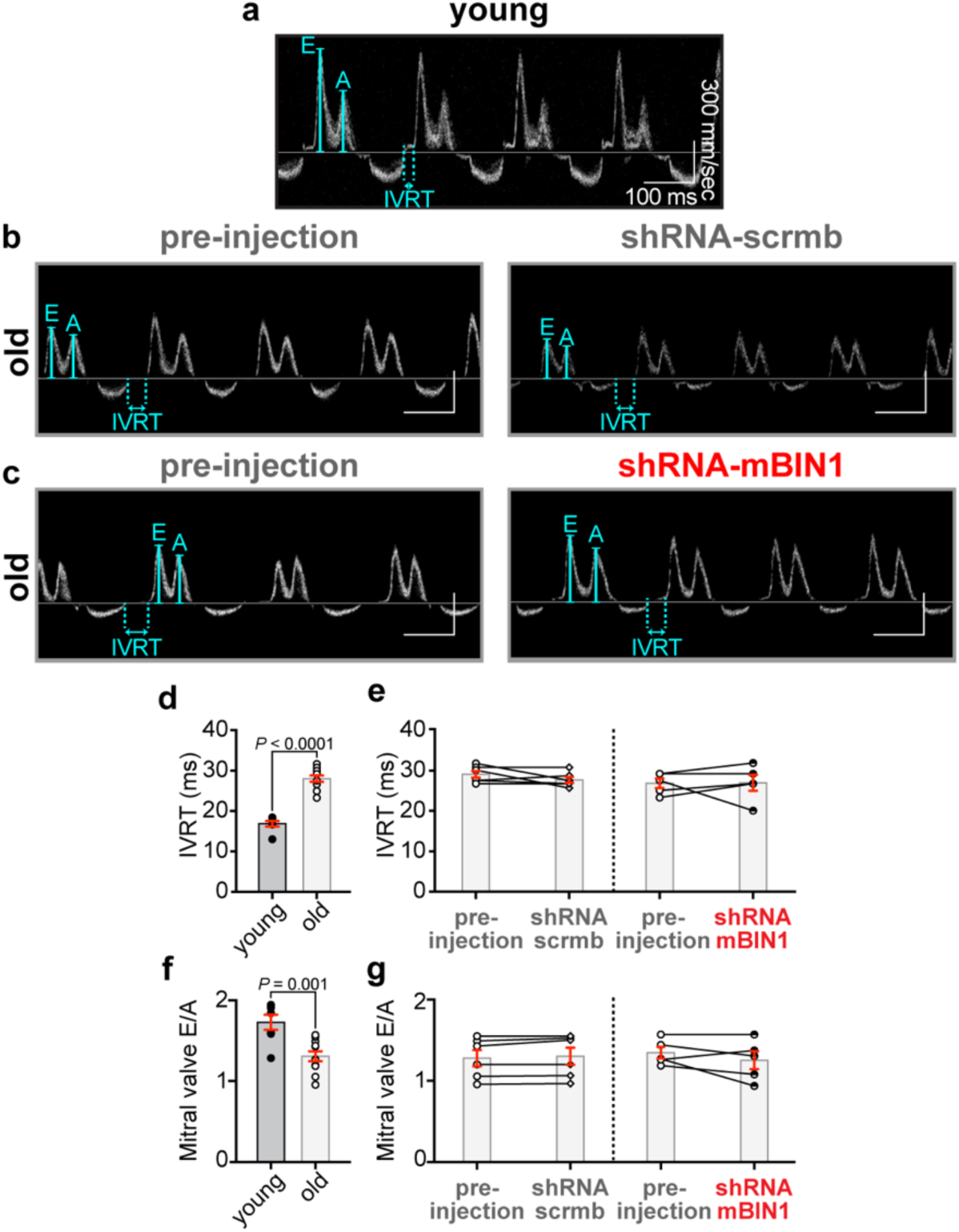
BIN1 knockdown does not recover age-related diastolic dysfunction. Representative pulsed-wave (PW) doppler images from unconscious young (**a**) and old mice before (left) and two-weeks after RO-injection (right) of shRNA-scrmb (**b**) and shRNA-mBIN1 (**c**). Summary dot-plots for young (*N* = 7) and old (*N* = 11) mice, and paired results before and after RO-injection of old mice with shRNA-scrmb (*N* = 6) and shRNA-mBIN1 (*N* = 5) for the following diastolic measurements are displayed: **d** and **e**, isovolumetric relaxation time (IVRT) and, **f** and **g,** mitral valve E/A ratio (MV E/A). Unpaired Student’s t-tests were performed on data displayed in d and f. Paired Student’s t-tests were performed on data displayed in e and g.

### Age-dependent decrease in myofilament Ca^2+^ sensitivity is rescued by BIN1 knockdown

Ca^2+^ sensitivity of the myofilaments can also influence systolic and diastolic function thus we examined the expression and phosphorylation state of two cardiac myofilament proteins that are known to impact Ca^2+^ sensitivity in a phosphorylation-dependent manner, specifically cardiac troponin-I (cTnI)^38, 39^ and cardiac myosin binding protein-C (cMyBP-C)^40, 41^. We probed for these myofilament proteins in heart lysates isolated from each cohort. We began with cTnI finding its total expression increased with aging, but relative expression of the phosphorylated form (pS23/24) reduced (Fig. 8a-c). While shRNA-mBIN1 treatment displayed a trend toward a more youthful pS23/24 expression, the trend did not become significant. Total expression of cMyBP-C was unaltered with aging but its phosphorylation at S273 was significantly decreased (Fig. 8d-f). shRNA-mBIN1 treatment restored youthful cMyBP-C pS273 expression levels. Our results support a model where reduced phosphorylation of cMyBP-C in old hearts contributes to deficits in contractile function while reduced cTnI phosphorylation favors longer relaxation and diastolic dysfunction. BIN1 knockdown appears to restore systolic but not diastolic function by improving phosphorylation of cMyBP-C but not cTnI.

**Figure 8.**
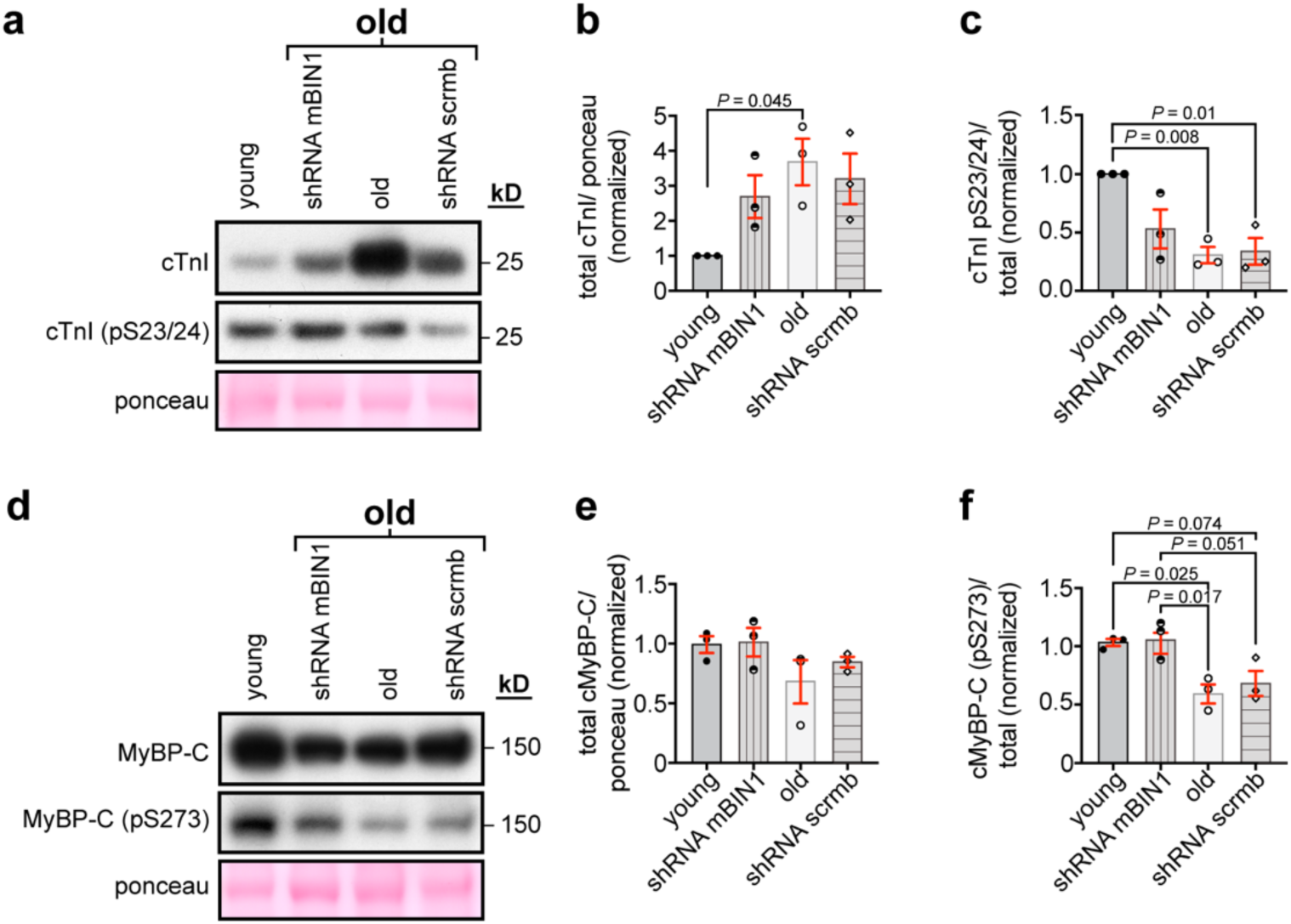
Myofilament Ca^2+^ sensitivity is altered in aging. **a**, western blot of total and pS23/24 cTnI expression in whole heart lysates from young, old, shRNA-mBIN1 or shRNA-scrmb transduced mice. Ponceau was used for normalization. **b**, histogram showing normalized total cTnI levels relative to young (*N* = 3, average of 3 replicates). **c**, histogram showing cTnI pS23/24 normalized to total cTnI relative to young (*N* = 3, average of 2-3 replicates). **d**, western blot of total and pS273 cMyBP-C expression in whole heart lysates from young, old, shRNA-mBIN1 or shRNA-scrmb transduced mice. Ponceau was used for normalization. **e**, histogram showing normalized total cMyBP-C levels relative to young (*N* = 3, average of 2-5 replicates). **f**, histogram showing cMyBP-C pS273 normalized to total cMyBP-C relative to young (*N* = 3, average of 2-5 replicates).

## Discussion

We report six novel findings that provide insight into the mechanistic basis of age-related cardiac dysfunction and reveal BIN1 as a new therapeutic target to rejuvenate the aging heart and extend cardiovascular health into old age: 1) patch clamp and Ca^2+^ transient recordings revealed an age-associated loss of cardiac *β*-adrenergic responsiveness measured by the functional augmentation of Ca_V_1.2 and RyR2 channel activity; 2) super-resolution imaging and PLA revealed an age-associated dysregulation of Ca_V_1.2 and RyR2 clustering plasticity, and juxta-positioning at dyads; 3) examination of endosomal pools and Ca_V_1.2 dynamics revealed ISO-stimulated Ca^2+^ channel mobility and recycling is impaired in aging due to trafficking deficits that cause endosomal traffic jams; 4) western blots and super-resolution imaging revealed that BIN1, important for t-tubule biogenesis^27^, and Ca_V_1.2, RyR2^18, 25, 26^ and sarcomere^42^ organization, is overexpressed in aging hearts; 5) shRNA-mediated BIN1 knockdown to youthful levels rejuvenated RyR2 clustering plasticity, basal Ca^2+^ transient amplitudes and their functional augmentation by *β-*AR-stimulation, and myofilament protein phosphorylation; and finally 6) *in vivo* echocardiography and Doppler imaging demonstrated that BIN1 knockdown in old mice restored youthful systolic (but not diastolic) function and substantially reversed the aging phenotype. Our knockdown approach using AAV9 packaged shRNA against BIN1 is rapid, occurring within 2 weeks of retro-orbital injection, and has translational potential as AAVs are approved for use in humans^43^.

Prior studies have shown that basal Ca_V_1.2 activity and *β*-AR-mediated *I*_Ca_ enhancement are altered with aging^44^. However, the effects on basal Ca_V_1.2 currents appear to vary depending on sex, species, and possibly even strain with various groups reporting enhanced^44–46^, unchanged^47^, or reduced^48–50^ currents in aged models. In our hands, myocytes from 24-month-old C57BL/6J mice had enhanced basal Ca_V_1.2 currents compared to young (3-month-old). Old myocytes also exhibited basally enhanced Ca^2+^ transient amplitudes. Prior findings on Ca^2+^ transient alterations in the aging myocardium range from basal amplitude augmentation^45, 51^ and faster decay kinetics^52^ to no or little reported change^50^, to basal reduction in CaT amplitude and slower rate of decay^46, 53^. This echoes the variability in age-associated alterations of *I*_Ca_ and reflects the complexity of aging. Importantly though, and in line with our results, there is agreement that the ability of *β*-AR-signaling to augment Ca^2+^ transient amplitude and tune EC-coupling and inotropy is diminished with aging^7, 8, 54–59^.

This study provides the first evidence linking age-associated *β*-AR hypo-responsivity to alterations in Ca^2+^ channel clustering and organization. Super-resolution microscopy studies of the nanoscale arrangement of Ca_V_1.2 and RyR2 have fueled the emerging concept that tunable insertion, recycling, and clustering of these channels physiologically during receptor signaling cascades or pathologically in disease, affects Ca^2+^ release efficiency and myocardial contractility^15, 16, 19, 20, 60–62^. Larger Ca_V_1.2 clusters facilitate enhanced Ca_V_1.2-Ca_V_1.2 physical interactions and cooperative gating that amplifies *I*_Ca_ ^16, 17, 63^ while larger RyR2 clusters favor enhanced Ca^2+^ spark frequency and larger Ca^2+^ transients^21^. We previously reported that acute *β*-AR activation in young ventricular myocytes promotes rapid, dynamic augmentation of sarcolemmal Ca_V_1.2 channel abundance as channels are mobilized from endosomal reservoirs via fast and slow recycling pathways^15, 16^. Here, super-resolution microscopy and dynamic channel tracking experiments demonstrated that this response is impaired with aging, such that Ca_V_1.2 are basally super-clustered to a similar extent to that seen in young ISO-stimulated myocytes. These results, correlated with the basally elevated *I*_Ca_ and impaired current augmentation with ISO, hint that the endosomal reservoir of Ca_V_1.2 cargo has already been expended and/or endosomal recycling pathways are impaired. Our data suggest the latter is true, revealing reduced ISO-induced insertions in old myocytes, an aggregation of channels on swollen endosomes, and a larger pool of static channels that appear stranded in the sarcolemma. Whether this pool of static channels reflects increased lifetime of Ca_V_1.2 in the membrane remains to be established. The main conclusion from this data is that the endosomal pathway and homeostatic regulation of Ca_V_1.2 channel expression at the sarcolemma is impaired with aging.

Deficiencies in recycling and endosomal enlargement are familiar sights in Alzheimer’s disease (AD) where endosomal enlargement is the first neuro-cytopathological hallmark of AD, visible even before neurofibrillary tangles or amyloid β plaques^23^. An imbalance in cargo trafficking into and out of endosomes is thought to result in endosome swelling and recycling pathway deficiencies termed endosomal traffic jams^24^. As mentioned earlier, alterations in BIN1 expression are thought to contribute to endosomal dysregulation in AD neurons. Thus, when we found BIN1 expression was significantly increased in aging, and appeared to assume a more endosomal localization pattern, we thought, since it was in the "right place at the right time" that knockdown of BIN1 would restore youthful endosomal trafficking and improve Ca_V_1.2 channel cargo mobilization, recovering *β-*AR responsivity and Ca_V_1.2 channel functional augmentation. That hypothesis was thoroughly disproven with results showing BIN1 knockdown did not in fact accomplish any of those things. Several other proteins have been implicated as sources or contributors to endosomal dysfunction and swelling in AD neurons as reviewed^24^. Future studies should investigate whether any of those proteins underlie age-associated endosomal dysfunction and impaired Ca_V_1.2 recycling in ventricular myocytes.

On the other side of the dyadic cleft, RyR2 also display enlarged basal clusters that fail to undergo augmentation with ISO in aging myocytes. Other groups have reported phosphorylation-stimulated clustering of RyR2^18, 19^ but this is the first report of disruption of this dynamic response with aging. Furthermore, it is the first to find that spatial proximity between Ca_V_1.2 and RyR2 increases with acute ISO in young myocytes. This suggests rapid dynamic formation of new dyads or expansion of existing dyads during ISO-stimulation. The implication here is that *β*-AR activation stimulates parallel Ca_V_1.2 and RyR2 cluster expansion on either side of the dyadic cleft to maximize the efficiency of Ca^2+^ release but that this additional tuning capacity or reserve is lost with aging and is apparently already expended at rest. Cardiac RyR2 trafficking and mobility is poorly understood and understudied but one report has linked RyR2 redistribution upon phosphorylation to BIN1, evidenced by reduced recruitment of phosphorylated RyR2 to dyadic regions in cardiac-specific *BIN1* knockout cardiomyocytes^18^. Interestingly, BIN1 knockdown in old mice had a more profound effect on RyR2 distribution, Ca^2+^ transient amplitude, and inotropy than it had on Ca_V_1.2. To that end, RyR2 nanoscale distribution was rejuvenated by BIN1 knockdown in old hearts but Ca_V_1.2 still exhibited the old super-clustered phenotype and heightened basal *I*_Ca_. So, while our initial rationale for knocking down BIN1 in the aging heart BIN1 was flawed, serendipitously we discovered that BIN1 knockdown has a rejuvenating effect on RyR2.

Echocardiography confirmed the benefits of BIN1 knockdown on the aging heart with parameters of systolic function including EF, FS, and CO all substantially improved after just a two-week knockdown of BIN1. EDV and ESV also declined to youthful levels reflecting the increased contractile function. In humans, resting systolic function is generally preserved with aging, however it becomes significantly impaired during exercise and acute stress^3, 64, 65^. The resting systolic dysfunction observed in the current study may reflect the difference between human and mouse heart’s basal autonomic balance at room temperature. Mice have relatively more sympathetic nervous system activity at rest than humans. With aging, autonomic balance becomes even more tilted toward sympathetic activity as parasympathetic input appears reduced^66, 67^. In this way, a resting mouse heart is not necessarily a fair comparison to a resting human heart and may instead be a better reflection of an exercising human heart. Thus, BIN1 knockdown may present a therapeutic option to enhance systolic function in aging humans during acute exercise and stress.

Diastolic dysfunction is a more universal finding with aging in both humans and mice^4, 68^ and our findings of a prolonged IVRT with aging and reduced E/A ratio agree with prior studies reporting impaired relaxation. However, BIN1 knockdown did not remedy diastolic dysfunction. This is perhaps expected as enhanced fibrosis^68^ with aging is one of the main causes of the enhanced stiffness that underlies diastolic issues in the aging heart, and there is no reason to suspect that BIN1 knockdown would improve fibrosis.

Reduced Ca^2+^ sensitivity of the myofilaments can also contribute to diastolic dysfunction. Enhanced phosphorylation of the thin myofilament protein cTnI by PKA at ser-23 and -24 accelerates myocardial relaxation by reducing myofilament Ca^2+^ sensitivity, favoring Ca^2+^ dissociation from troponin-C, and leading to faster actin-myosin detachment^38, 39^. Our results indicate that aging hearts display significantly lower levels of cTnI phosphorylation favoring slower relaxation. This deficit was not improved or corrected by BIN1 knockdown and diastolic dysfunction prevailed. Given the reciprocal relationship between cTnI phosphorylation and myofilament Ca^2+^ sensitivity, one would predict that the age-associated reduction in cTnI pS23/24 levels would result in increased Ca^2+^ sensitivity in old mice compared to young, leading to increased Ca^2+^-dependent force generation. However, our *in vitro* finding of larger amplitude Ca^2+^ transients in old myocytes accompanied by reduced *in vivo* contractile function in old mice suggests quite the opposite. Specifically, our functional results suggest an age-associated reduction in the Ca^2+^ sensitivity of the myofilaments that is remedied by BIN1 knockdown. Thus, we broadened our search to examine cMyBP-C, another myofilament protein. Decreased phosphorylation of cMyBP-C causes decreases myofilament Ca^2+^ sensitivity^41^ and contractile dysfunction in heart failure patients^40^. Furthermore, in the dephosphorylated state, cMyBP-c inhibits actin-myosin interactions leading to reduced force development^41^. We found old mouse hearts had lower cMyBP-C pS273^69^ expression levels than young hearts. A similar age-associated decline in cMyBP-C phosphorylation has been reported by others and found to correlate with deterioration of cardiac function^70^. Furthermore, phosphorylation-mimetic cMyBP-C mutant mice exhibit preserved cardiac function and enhanced longevity suggesting phosphorylation of cMyBP-C is cardioprotective^70^. In a novel finding, we report that BIN1 knockdown rescues cMyBP-C pS273 phosphorylation levels which potentially explains the improved contractile function and Ca^2+^ sensitivity in shRNA-mBIN1 transduced mice. But how can one link BIN1 expression levels to sarcomeric proteins? While we do not yet have the full mechanistic picture of how this occurs, there is evidence that BIN1 binds to and influences sarcomeric proteins. For example, in skeletal muscle, BIN1 is said to act as an essential adaptor protein and scaffold that promotes mature sarcomere assembly^42^. Increased BIN1 expression is also known to promote enhanced expression of myosin heavy chain^71^ and to associate with and stabilize actin filaments^72^. Future studies should examine the mechanism of BIN1 regulatory influences over cMyBP-C but it is an intriguing idea first proposed by Fernando et al in 2009^42^, that a single protein (BIN1) could orchestrate sarcolemmal and sarcomeric architecture and regulation.

It is important to mention that although our study focused on the effects of BIN1 upregulation with aging, prior work has revealed that BIN1 *downregulation* is associated with heart failure^26, 29^ and that exogenous supplementation of a cardiac specific isoform of BIN1 can improve cardiac function in mice with pressure overload-induced heart failure^73^. It is possible that BIN1 levels must be tightly regulated and that any deviation from a homeostatic normal level could be damaging. In Alzheimer’s disease such a model has been presented where too much or too little BIN1 expression are both associated with enhanced risk of disease^35, 74^.

In summary, we find that BIN1 knockdown restores youthful Ca^2+^ transient amplitude, *β*-AR mediated transient augmentation, and *in vivo* systolic function in the aging myocardium. This study sets the groundwork for future work and raises the hope that a similar approach could improve cardiac function, restore exercise tolerance, and increase capacity to endure acute stress in elderly patients.

## Acknowledgments

We extend our thanks to Dr. Sakthivel Sadayappan for the generous gift of the total cMyBP-C antibody and the cMyBP-C Ser-273 phospho-site specific antibody^69^. We thank Dr. L. Fernando Santana and Heather Spooner for reading and providing critical suggestions on our manuscript.

## Sources of Funding

This work was supported by NIH R01AG063796 and R01HL159304 to R.E.D.; R01GM127513 and RF1NS131379 to E.J.D.; by the NIGMS funded Pharmacology Training Program T32 GM099608 and later by an AHA Predoctoral Fellowship 827909 to T.L.V., by a Postdoctoral Fellowship from NIH T32 Training Grant in Basic & Translational Cardiovascular Science T32 HL086350 and NIH F32 HL149288 to P.N.T.; and by NIH R01HL085727, R01HL085844, R01HL137228, S10OD010389 Core Equipment Grant, and VA Merit Review Grant I01 BX000576 and I01 CX001490 to N.C. Graphical Abstract was generated using Biorender.com.

## Disclosures

None.

## Notes

### Competing Interest Statement

The authors have declared no competing interest.

### Summary of Updates

New data has been added including echocardiography and doppler imaging.

