## Supplemental Material for "BIN1 knockdown rescues systolic dysfunction in the aging heart"

### Supplementary Material

### Detailed Methods

#### Ethical Approval

University of California Davis Institutional Animal Care and Use Committee (IACUC) approved all procedures involving mice which were conducted in accordance with the Guide for the Care and Use of Laboratory Animals[1]. Mice received food and water *ad libitum*.

#### Isolation of mouse ventricular myocytes

All experiments were performed on hearts from male mice, except in Figure 5a-b where male and female hearts were used. Hearts from 3-5-month-old (referred to as "young"; The Jackson Laboratory, Sacramento, CA, USA) and 21-25-month-old (referred to as "old"; National Institute on Aging Aged Rodent Colony) C57BL/6J mice were surgically removed after intraperitoneal injection of pentobarbital solution (> 100 mg/kg; B euthanasia-D Special; Merck & Co., Inc., Rahway, NJ, USA). Upon removal, hearts were repeatedly plunged into chilled digestion buffer (130 mM NaCl, 5 mM KCl, 3 mM Na-pyruvate, 25 mM HEPES, 0.5 mM $MgCl_2$, 0.33 mM $NaH_2PO_4$, 22 mM glucose, and 150 µM EGTA) to encourage replacement of blood in the chambers and ventricular myocytes were isolated using the Langendorff technique as previously described[2, 3]. Briefly, aortic cannulation was performed and used to retrogradely perfuse the heart with 37°C digestion buffer supplemented with 50 µM $CaCl_2$ (Thermo Fisher Scientific, Rockford, IL, USA), 0.04 mg/ml protease (Sigma-Aldrich, Inc., St. Louis, MO, USA) and 1.4 mg/ml type 2 collagenase (Worthington Biochemical, Lakewood, NJ, USA). The decision that digestion was complete was made by monitoring the color, shape, and texture of the perfused heart, removing it from the Langendorff system when it appeared pale, flaccid, and soft to the touch. Ventricles were then cut away from the rest of the heart, chopped into smaller pieces, and myocytes were dissociated by gently pipetting the pieces in 37°C digestion buffer supplemented with

0.96 mg/mL type 2 collagenase, 0.04 mg/mL protease, 100 μM $CaCl_2$ and 10 mg/ml BSA (Sigma-Aldrich). After 2 min centrifugation at 300 rpm, the supernatant was discarded, and cells were resuspended in a wash buffer consisting of the digestion buffer supplemented with 10 mg/ml BSA and 250 μM $CaCl_2$. A final 2 min centrifugation step was performed, the supernatant wash buffer was aspirated away and the cells were resuspended in the appropriate solution for the planned experiments.

Strict standards for isolation and cell quality were adhered to. Accordingly, cells were considered "healthy" and pursued for experiments when they fulfilled the following criteria:

(1) Cell isolation: at least 75 % cell survival was required to consider an isolation successful and to proceed with further evaluation and experiments.

(2) Cell morphology: "rod-shaped"/"brick-shaped" cells with clear striations, no blebbing, and no bunching at the cell ends.

(3) Cell stability: cells were required to be quiescent in physiological (1.8 mM $Ca^{2+}$-containing) solution, with no spontaneous contractions.

(4) When electrophysiology was performed, cells were pursued if they were found capable of holding stable seals with no more than 200 pA leak (but usually < 50 pA) for at least 10 mins.

**Whole-cell patch clamp electrophysiology**

Whole-cell patch clamp was performed on freshly isolated ventricular myocytes to record $Ca_V1.2$ channel currents ($I_{Ca}$). Borosilicate glass pipettes (Sutter instrument, Novato, CA, USA) were fire polished to 1-3 MΩ resistance, and filled with an internal solution containing 87 mM Cs-aspartate, 20 mM CsCl, 1mM $MgCl_2$, 10 mM HEPES, 10 mM EGTA and 5mM MgATP (pH adjusted to 7.2 with CsOH). The MgATP was added into the solution on the day of experiments.

Initially, myocytes were perfused with an external Tyrode's solution containing 140 mM NaCl, 5 mM KCl, 10 mM HEPES, 10 mM Glucose, 1 mM $MgCl_2$ and 2 mM $CaCl_2$ (pH adjusted to 7.4 with NaOH). Once a whole-cell configuration was obtained, perfusion was switched to a solution containing 5 mM CsCl, 10 mM HEPES, 10 mM Glucose, 140 mM NMDG, 1 mM $MgCl_2$ and 2 mM $CaCl_2$ (pH adjusted to 7.3 with HCl).

To minimize the effects of $I_{Ca}$ rundown, cells were held at a holding potential of -80 mV for 5 mins prior to the recording of control currents. Next, cells were held at -80 mV and, to inactivate $Na^+$ currents, stepped to -40 mV for 100 ms, followed by 300 ms steps to voltages ranging from -60 to +90 mV. Recordings were obtained in control conditions and after 2-3 mins of perfusion with 100 nM isoproterenol (ISO; Sigma-Aldrich) in the external bath solution. Currents were sampled at a frequency of 10 kHz, and low-pass-filtered at 2 kHz using an Axopatch 200B amplifier (Molecular Devices, Sunnyvale, CA, USA), digitized using a Digidata 1550B plus Humsilencer (Molecular Devices) and acquired using pClamp (Molecular Devices). Analysis was performed using Clampfit software (Molecular Devices). Membrane potentials were corrected for a liquid junction potential of -10 mV.

**Calcium transient recordings**

Freshly isolated cardiomyocytes were loaded with 10 µM Fluo4-AM or 10 µM Rhod-2 AM (Thermo Fisher Scientific) for 20 min in the dark. After loading, myocytes were liberated from excess indicator via centrifugation for 2 min at 300 rpm and were subsequently resuspended in Tyrode's solution where they de-esterified for a further 20 min prior to commencement of experiments.

Myocytes were paced at 1 Hz using a 12V square-wave stimulus evoked by a Myopacer electric field stimulator (IonOptix, LLC., Westwood, MA) and the resulting $Ca^{2+}$ transients were visualized by exciting Fluo-4 (with 488-nm laser light) or Rhod-2 (with 594-nm laser light) and capturing a sequence of line-scans across the length of the cell using a Zeiss LSM 880 line-scanning confocal microscope (Carl Zeiss Microscopy, LLC., White Plains, NY, USA) equipped with a Plan-Apochromat 63×/1.40 N.A. oil immersion objective. Cells were continuously perfused with Tyrode's solution until steady state was achieved when control transients were recorded. The same cell was then perfused with 100 nM ISO-containing Tyrode's for 3 min before capturing a second group of transients to assess the ISO-induced effects.

Fluo-4 and Rhod-2 fluorescent signals were converted into intracellular $Ca^{2+}$ concentration using the pseudo-ratiometric approach[4] and the equation: $[Ca^{2+}]_i = K_d(F/F_0)/(K_d/[Ca^{2+}]_{i\text{-rest}}+1-F/F_0)$ as previously described[5, 6].

**Single Molecule Localization Microscopy**

Coverslips (#1.5; VWR, Radnor, PA, USA) were sonicated for 20 mins in 1M NaOH to remove any contaminants, followed by several washes in de-ionized water and stored in 70% ethanol until use. Freshly isolated ventricular myocytes were plated onto poly-$_L$-lysine (0.01%; Sigma-Aldrich); and laminin (20 µg $ml^{-1}$; Life Technologies, Carlsbad, CA, USA) coated coverslips and left for 45 mins in a 37°C incubator to adhere. Adhered cells were then treated for 8 min with either PBS (control) or 100 nM ISO in PBS. The cells were then fixed and permeabilized in 100% ice-cold methanol (Thermo Fisher Scientific) for 5 min at -20°C, followed by several washing steps in PBS. Fixed cells were blocked and permeabilized for 1 hr at room temperature in 20% SEA BLOCK blocking buffer (Thermo Fisher Scientific) and 0.25% v/v Triton X-100 (Sigma-Aldrich) in PBS.

Cells were incubated overnight at 4°C in rabbit polyclonal IgG anti-$Ca_V$1.2 (CACNA1C, ACC-003, Alomone Labs, Jerusalem, Israel; 1:300 dilution) or mouse monoclonal IgG1 anti-RyR2 (C3-33, MA3-916, Invitrogen, Waltham, MA, USA; 1:50 dilution) in blocking buffer. Excess primary antibodies were washed off with PBS before subsequent incubation in relevant secondary antibodies [Alexa Fluor 647-conjugated donkey anti-rabbit or Alexa Fluor 647-conjugated donkey anti-mouse IgG1 (Invitrogen; 1:1000 dilution in blocking buffer)] for 1 hr at room temperature. After multiple washes in PBS, coverslips were mounted onto glass depression slides (neoLab, Heidelberg, Germany) with a cysteamine (MEA)-catalase/glucose/glucose oxidase (GLOX) imaging buffer containing TN buffer (50 mM Tris pH 8.0, 10 mM NaCl), a GLOX oxygen scavenging system (0.56 mg $mL^{-1}$ glucose oxidase, 34 μg $mL^{-1}$ catalase, 10% w/v glucose) and 100 mM MEA.

To exclude oxygen and hold coverslips in place, Twinsil silicone-glue (Picodent, Wipperfürth, Germany) and aluminum tape (T204-1.0-AT205; Thorlabs Inc., Newton, NJ, USA) were used. Fixed cells were imaged on a Leica DMi8 microscope (Leica Microsystems, Wetzlar, Germany) in HiLo TIRF mode, using a HC PL APO 160x 1.43 oil CORR GSD objective (Leica Microsystems). Ground state depletion was performed, and dye-blinking was elicited using a 638 nm/150 mW laser. Photon emission was detected with a Hamamatsu Flash 4.0 camera. Raw blinking images were collected with an exposure time of 10 ms for 50,000 frames using LAS X Life Science Software (Leica Microsystems). Localization maps with 10 nm pixel size were generated using a detection threshold of 30 and camera gain factor of 2.2. Mean $Ca_V1.2$ and RyR2 cluster areas were quantified using the "analyze particles" function in ImageJ/FIJI as described previously[7].

**Proximity Ligation Assay**

Coverslip adhered ventricular myocytes were treated for 8 min with PBS (control) or 100 nM ISO. At the conclusion of the treatment period, cells were fixed for 20 mins in 4% paraformaldehyde (Electron Microscopy Sciences, Hatfield, PA, USA) in PBS at room temperature. Excess aldehyde groups were quenched with a subsequent 15 min treatment with 100 mM glycine (Sigma-Aldrich) in PBS, followed by 2x 3 min washes with PBS. Cells were next permeabilized with 0.1% Triton X-100 in PBS for 20 min and were then thoroughly washed before undergoing a 1hr room temperature blocking step in 20% SEA BLOCK and 0.25% v/v Triton X-100 in PBS.

Cells were incubated overnight at 4°C in primary antibody solutions diluted in blocking buffer. Primaries utilized were affinity-purified rabbit polyclonal antibody Cav1.2 II-III (RRID:AB_2922674, courtesy of the Trimmer Lab, UC Davis, Davis, CA, USA; 1:100) and mouse monoclonal IgG1 anti-RyR2 (C3-33, MA3-916, Invitrogen; 1:50). The manufacturer's instructions were then followed for the Duolink PLA Fluorescence protocol (Sigma-Aldrich). This involved incubating samples with PLA probes (anti-mouse MINUS and anti-rabbit PLUS) for 1hr at 37°C, followed by a ligation step (30 min at 37°C), an amplification step (100 min at 37°C) and final washes in Duolink buffer B. Coverslips were mounted on a microscope slide with DAPI-containing Duolink *In Situ* mounting medium (Sigma-Aldrich) and cells were examined using a Zeiss LSM 880 super-resolution microscope equipped with an Airyscan detector and a Plan-Apochromat 63x/1.40 oil DIC M27 objective. Images were acquired using Zen software (Carl Zeiss Microscopy, LLC). Maximum intensity z-projections (from 0.5 µm slice intervals) were analyzed using the "analyze particles" function in ImageJ/FIJI to quantify the puncta per area for each cell.

**Total Internal Reflection Fluorescence (TIRF) imaging of live transduced cardiomyocytes**

Isoflurane-anesthetized mice were retro-orbitally injected with AAV9-$Ca_v\beta_{2a}$-paGFP at a concentration of $4 \times 10^{12}$ vg/ml 4-6 weeks before the planned experiment when transduced myocytes were isolated and plated onto poly-$_{\text{L}}$-lysine coated coverslips as described above. Photoactivation of the transduced $Ca_v\beta_{2a}$-paGFP was achieved by illumination with 405 nm LED light and the photoactivated GFP was subsequently excited with 488 nm laser light at a TIRF penetration depth of 153 nm on an Olympus IX83 inverted microscope equipped with a Cell-TIRF MITICO and a 60x/1.49 N.A. TIRF objective lens and an iXon Ultra 888 back thinned EM-CCD camera (Andor, Belfast, Northern Ireland). Images were acquired at 10.34 FPS. After a control period of 300 frames with Tyrode's solution perfusion, cells were stimulated with 100 nM ISO for 3 min (1861 frames).

ImageJ/FIJI was used to analyze and quantify $Ca_v\beta_{2a}$-paGFP populations. Images were bleach corrected before application of a 20-pixel rolling ball background subtraction and a 10-frame moving average. Maximum intensity z-projections of the first 300 frames of each experiment were used to represent the control period and the final 300 frames of the ISO perfusion were used to represent the ISO-stimulated period. These control and ISO period z-projections were then subjected to thresholding to binarize them and image math was performed to visualize and quantify the following channel populations, those that were: i) inserted/recycled during the ISO period (ISO – Ctrl), ii) endocytosed during the course of the experiment (Ctrl – ISO); and iii) those that remained static throughout (Ctrl * ISO) as described previously[2].

**Fixed cell immunostaining and Airyscan microscopy**

Freshly isolated ventricular mouse myocytes were plated onto poly-$_{L}$-lysine and laminin-coated coverslips and subjected to control (PBS) or 8 min ISO-stimulation in a 37°C incubator prior to fixation with 4% paraformaldehyde solution with a cytoskeletal preserving buffer (PEM) containing 80 mM PIPES, 5 mM EGTA, and 2 mM $MgCl_2$ for 10 min. After PBS rinse, cells were permeabilized for 10 mins in 0.5% Triton-X 100 (Sigma-Aldrich) and, after several rinses with PBS, blocked with 20% SEA Block (Thermo Fisher Scientific) and 0.25% v/v Triton X-100 for 1 hr at room temperature. Primary antibodies diluted in blocking solution for overnight incubation were: rabbit polyclonal IgG anti-$Ca_v$1.2 (ACC-003, Alomone Labs; 1:300), mouse monoclonal anti-EEA1 IgG1 (ab70521, Abcam, Waltham, MA, USA; 1:250), mouse monoclonal $IgG_1$ anti-Rab11a (sc-166523, Santa Cruz Biotechnology Inc., Dallas, TX, USA; 1:250;), mouse monoclonal $IgG_1$ anti-Rab7 (sc-376362, Santa Cruz; 1:250) and mouse monoclonal $IgG_1$ anti-Bin1, clone 2F11 (NBP2-21675, Novus Biologicals, Littleton, CO, USA; 1:125). Cells were then washed with PBS and incubated for 45 min at RT with the relevant Alexa Fluor conjugated secondary antibodies (Life Technologies; 1:1000). Secondaries utilized included Alexa Fluor 488 goat anti-rabbit, Alexa Fluor 647 goat anti-mouse $IgG_1$ and Alexa Fluor 488 goat-anti mouse $IgG_1$ (Invitrogen; 1:1000). Coverslips were mounted onto glass slides and examined on a Zeiss LSM 880 super-resolution microscope equipped with an Airyscan detector and a Plan-Apochromat 63×/1.40 oil DIC M27 objective and Zen software.

**Western blot**

Mice were euthanized as described, then whole hearts were collected and flash-frozen in liquid nitrogen. Frozen hearts were then homogenized in RIPA buffer (R26200, Research Products International, Mount Prospect, IL USA) containing protease inhibitor, 4μM microcystin (Insolution microcystin-LR, Thermo Fisher Scientific, Rockford, IL, USA) and 1mM NaF using tubes pre-filled with beads (Next Advance, Inc., Troy, NY, USA) for centrifugation at 4°C for 20 min. The supernatant was then extracted, and protein concentration was determined by BCA assay. All protein samples were denatured in SDS sample buffer (Bolt SDS Sample Buffer, Invitrogen) at 70°C for at least 10 mins. The protein was fractionated by SDS-PAGE 4-12% gradient acrylamide gels and transferred onto polyvinylidene difluoride membranes (PVDF; Bio-Rad Laboratories, Hercules, CA, USA) using transfer buffer (95.9 mM glycine, 12.5 mM Tris, 150 ml MeOH, up to 1 L $H_2$0) at 50 V for 10 hrs. PVDF membranes were ponceau stained and imaged using a photo scanner (Epson Perfection V600 photo). The membrane was cut into separate bands of interest and incubated in a blocking buffer (BB) consisting of 150 mM NaCl, 10 mM Tris-HCl, pH 7.4 (TBS) with 0.2 % Tween (TBST) and 5 % milk for 1 hr at RT. Next, they were incubated with primary antibodies in BB overnight at 4°C. BIN1 was detected by anti-BIN1 (2F11, Novus Biologicals, or 99D, Sigma-Aldrich) each at 1:400. Total cTNI was detected by anti-troponin I, (TI-4, 1:100; TI-4 was deposited by Schiaffino, S to the Developmental Studies Hybridoma Bank, created by the NICHD of the NIH and maintained at The University of Iowa, Department of Biology, Iowa City, IA 52242). Phosphorylated cTnI (pS23/24) was detected by anti-phospho-troponin I (Cardiac) (Ser23/24) (Cell Signaling Technology 4004, 1:1000). Total cMyBP-C was detected by anti-cMyBP-C 2-14 (kindly provided by Sakthivel Sadayappan, University of Cincinnati, 1:10,000). Phosphorylated cMyBP-C was detected by anti-Ser 273 (kindly provided by Sakthivel Sadayappan, University of Cincinnati, 1:2500).

Membranes were washed for at least 30 min with at least five exchanges of BB. Membranes were subsequently incubated with horseradish peroxidase-conjugated secondary antibodies: anti-Mouse IgG (H+L)-HRP Conjugate (Biorad 1706516) and monoclonal mouse anti-rabbit IgG light chain specific HRP conjugate (Jackson ImmunoResearch 211-032-171) both diluted in BB at 1:10,000, rocking for 1 hour at RT. Secondary antibodies were then washed with TBST for 90 min with at least 6 exchanges of solution. The membranes were developed using chemiluminescent reagents (Immobilon Classico, Sigma-Aldrich; Prometheus ProSignal Femto, Genesee Scientific, El Cajon, CA, USA) on autoradiography film (Amersham Hyperfilm ECL, Cytivia, Global Life Sciences Solutions USA, LLC., Marlborough, MA, USA) using a film developer to quantify protein expression. Multiple scans were taken over varying time periods to ensure signals were in the linear range and bands were non-saturated. Films were scanned using the photo scanner and immunoreactive bands were quantified using densitometry in Adobe Photoshop, normalizing to protein loading using the Ponceau scan. Note, total protein is the only acceptable normalization standard in aging studies since the protein levels of many, if not all, the standard 'housekeepers' change with aging[8, 9]. Individual lysates (biological replicates) were probed 2-5 times on separately run gels constituting technical replicates.

**Adeno-associated virus (AAV9) short hairpin RNA (shRNA) for BIN1 knockdown**

AAV9-GFP-U6-m-Bin1-shRNA (shRNA-mBIN1) and AAV9-GFP-scrmb-shRNA (shRNA-scrmb) were purchased from Vector Biolabs (Malvern, PA, USA). For BIN1 knockdown, 22–25-month-old mice were anesthetized with vaporized isoflurane and retro-orbitally injected with the shRNA-mBIN1 or shRNA-scrmb at a concentration of $5 \times 10^{11}$ vg/ml two weeks before experiments when animals were euthanized, and myocytes isolated, or hearts harvested for biochemistry.

**Echocardiography**

Echocardiography was performed using a Vevo 2100 imaging system (VisualSonics, Fujifilm, Toronto, ON, Canada) and a MS 550D probe (22-55 MHz). Systolic function from conscious and systolic and diastolic function from unconscious (anesthetized with 2 % isofluorane) young and old mice was evaluated using M-mode echocardiography and pulsed-wave Doppler. Two-dimensional measurements were used to extract fractional shortening (FS), ejection fraction (EF), left ventricular (LV) mass, heart rate (HR), end diastolic volume (EDV), end systolic volume (ESV), cardiac output (C), mitral valve E/A wave ratio (MV E/A) and isovolumetric relaxation time (IVRT) values as described previously[10, 11]. Intraperitoneal injection of 0.1 mg/kg ISO diluted in saline was used on unconscious mice to measure β-AR response.

For paired BIN1 knockdown results, echocardiography was performed on WT old mice prior to and two weeks after retro-orbital injections of AAV9-shRNA-scramble or AAV9-shRNA-mBIN1.

**Statistical analysis**

*N* represents the number of animals and *n* represents the number of cells. Data are reported as mean ± SEM. GraphPad Prism software (GraphPad Software Inc., La Jolla, CA, USA) was used to graph and compare datasets using paired or unpaired Student's t-tests, one-way ANOVAs, or two-way ANOVAs with Tukey's multiple comparison post-hoc tests as stated in the specific figure legends. $P < 0.05$ was considered statistically significant.

### Supplementary Figures and Figure Legends

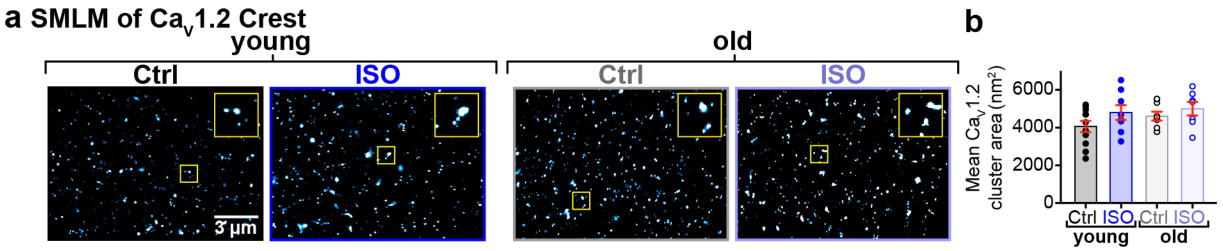


**Supplementary Figure 1. $Ca_V1.2$ clustering at the sarcolemmal crest is unaltered by aging.** **a**, SMLM localization maps showing $Ca_V1.2$ channel localization and distribution in the sarcolemmal crest of young and old ventricular myocytes with or without ISO-stimulation. Yellow boxes indicate the location of the regions of interest magnified in the top right of each image. **b**, dot-plots summarizing the mean $Ca_V1.2$ cluster areas in young (control: $N$ = 4, $n$ = 11; ISO: $N$ = 5, $n$ = 8) and old (control: $N$ = 3, $n$ = 9; ISO: $N$ = 3, $n$ = 7) myocytes. Data were analyzed using two-way ANOVAs with post-hoc Tukey's multiple comparisons tests.

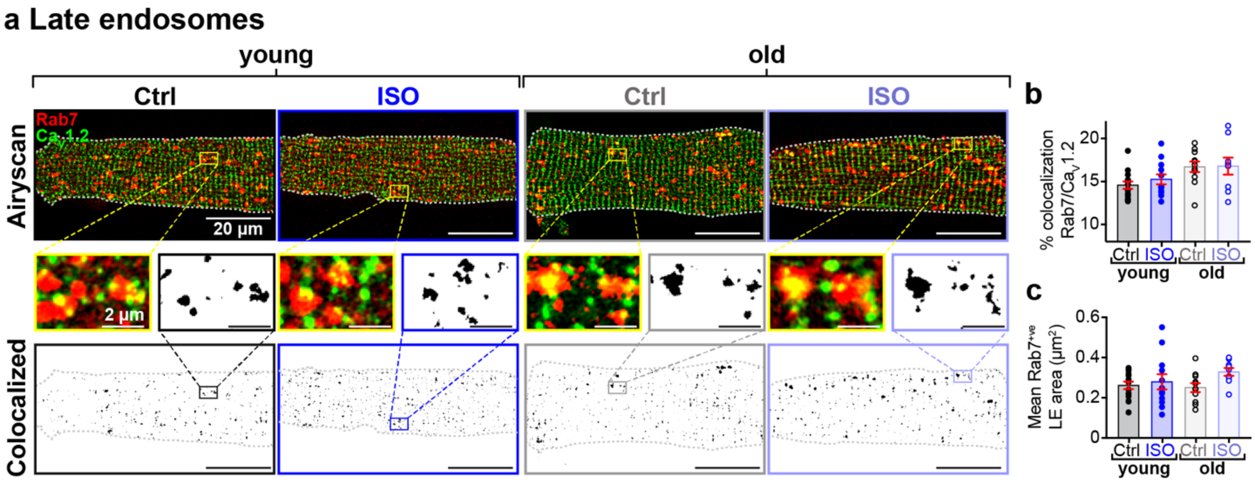


**Supplementary Figure 2. Rab7 positive late endosome $Ca_V1.2$ content and size are not altered in aging myocytes.** **a**, Airyscan super-resolution images of Rab7 and $Ca_V1.2$ immunostained young and old myocytes under control and ISO-stimulated conditions. *Bottom:* Binary colocalization map showing overlapped expression. **b**, dot-plots summarizing % colocalization between Rab7 and $Ca_V1.2$, and **c**, mean area of Rab7 positive endosomes in young (control: $N$ = 3, $n$ = 14; ISO: $N$ = 3, $n$ = 12) and old (control: $N$ = 3, $n$ = 11; ISO: $N$ = 3, $n$ = 9) myocytes. Statistical analyses were performed with two-way ANOVAs and post-hoc Tukey's multiple comparisons tests.

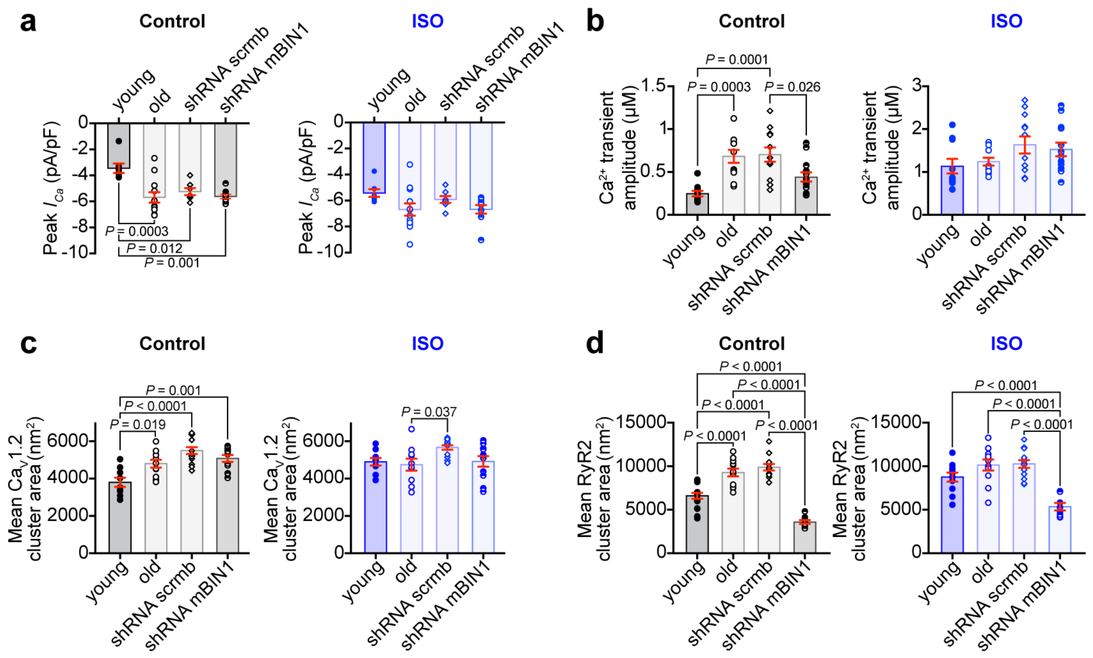


**Supplementary Figure 3. Further characterization of BIN1 knockdown in old mice. a**, dot-plots showing peak $I_{Ca}$ in control (left) and ISO (right) conditions for young, old, shRNA-scrmb ($N$ = 3, $n$ = 7) and shRNA-mBIN1 ($N$ = 3, $n$ = 9) transduced old myocytes. Old and young data points are reproduced from Fig. 1c. **b**, dot-plots summarizing $Ca^{2+}$ transient amplitude in control (left) and ISO (right) conditions from young, old, shRNA-scrmb ($N$ = 3, $n$ = 12) and shRNA-mBIN1 ($N$ = 5, $n$ = 14) transduced old myocytes. Old and young data points are reproduced from Fig. 1f. **c**, dot-plots summarizing the mean $Ca_V1.2$ channel cluster areas in control (left) and ISO (right) conditions for young, old, shRNA-scrmb (control: $N$ = 3, $n$ = 14; ISO: $N$ = 3, $n$ = 13) and shRNA-mBIN1 (control: $N$ = 3, $n$ = 12; ISO: $N$ = 3, $n$ = 11) transduced old myocytes. Old and young data points are reproduced from Fig. 2b. **d**, dot-plots summarizing the mean RyR2 channel cluster areas in control (left) and ISO (right) conditions for young, old, shRNA-scrmb (control: $N$ = 3, $n$ = 12; ISO: $N$ = 3, $n$ = 13) and shRNA-mBIN1 (control: $N$ = 3, n = 9; ISO: $N$ = 3, $n$ = 8) transduced old myocytes. Old and young data points are reproduced from Fig. 2d. Statistical analysis was performed using one-way ANOVAs with post-hoc Tukey's multiple comparison tests.

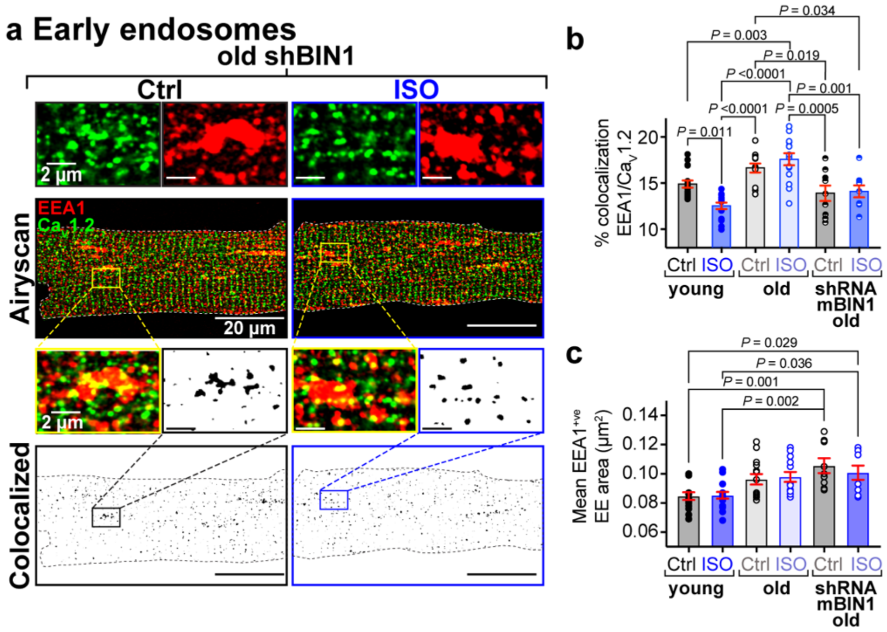


**Supplementary Figure 4. EEA1 positive early endosomes are not altered with BIN1 knockdown. a**, Airyscan super-resolution images of EEA1 and Ca$_V$1.2 immunostained shRNA-mBIN1 transduced old myocytes under control and ISO-stimulated conditions. *Bottom:* Binary colocalization map showing overlapped expression. **b**, dot-plots summarizing % colocalization between EEA1 and Ca$_V$1.2, and **c**, mean area of EEA1 positive endosomes in young, old and shRNA-mBIN1 (control: $N$ = 3, $n$ = 9; ISO: $N$ = 3, $n$ = 9 ) transduced old myocytes. Old and young data points are reproduced from Fig. 4b and c. Statistical analyses in b and c were performed using two-way ANOVAs and post-hoc Tukey's multiple comparisons tests.

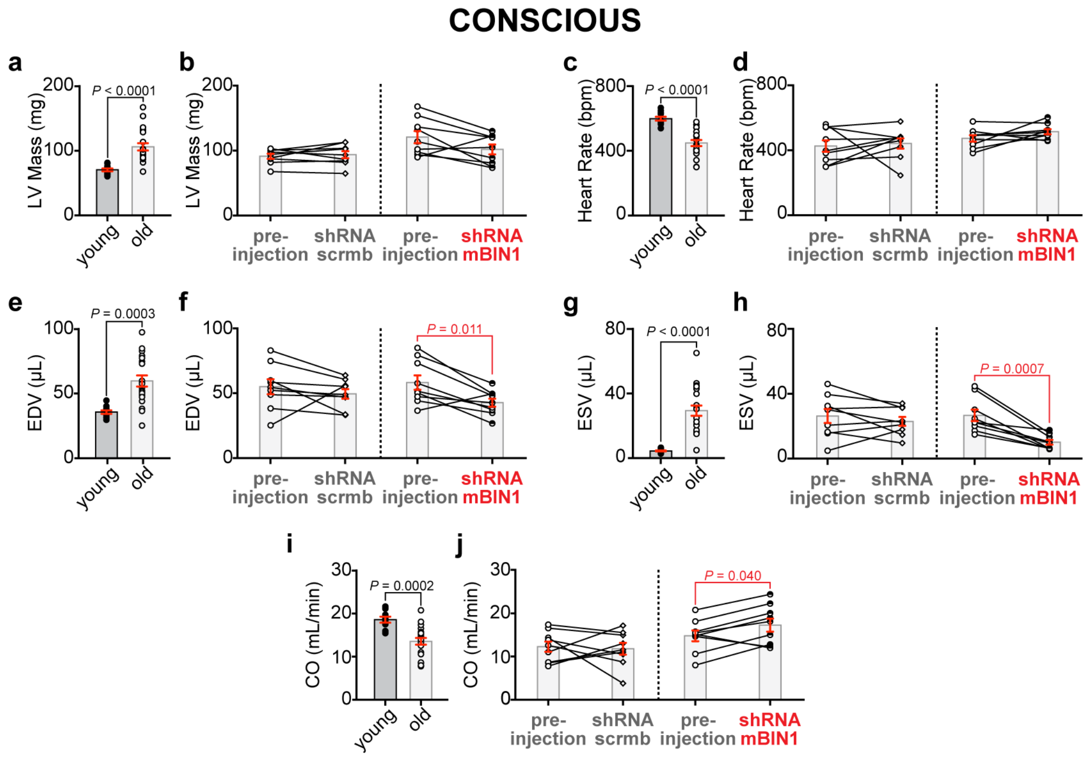


**Supplementary Figure 5. Additional data from conscious M-mode echocardiography.** Summary dot-plots for conscious young (*N* = 11) and old (*N* = 20) mice, and paired results before and after RO-injection of old mice with shRNA-scrmb (*N* = 9) and shRNA-mBIN1 (*N* = 9) for the following measurements are displayed: **a** and **b**, left ventricular (LV) mass, **c** and **d**, heart rate (HR), **e** and **f**, end diastolic volume (EDV), **g** and **h**, end systolic volume (ESV) and, **i** and **j**, cardiac output (CO). Unpaired Student's t-tests were performed on data displayed in a, c, e, g, and i. Paired Student's t-tests were performed on data displayed in b, d, f, h, and j.

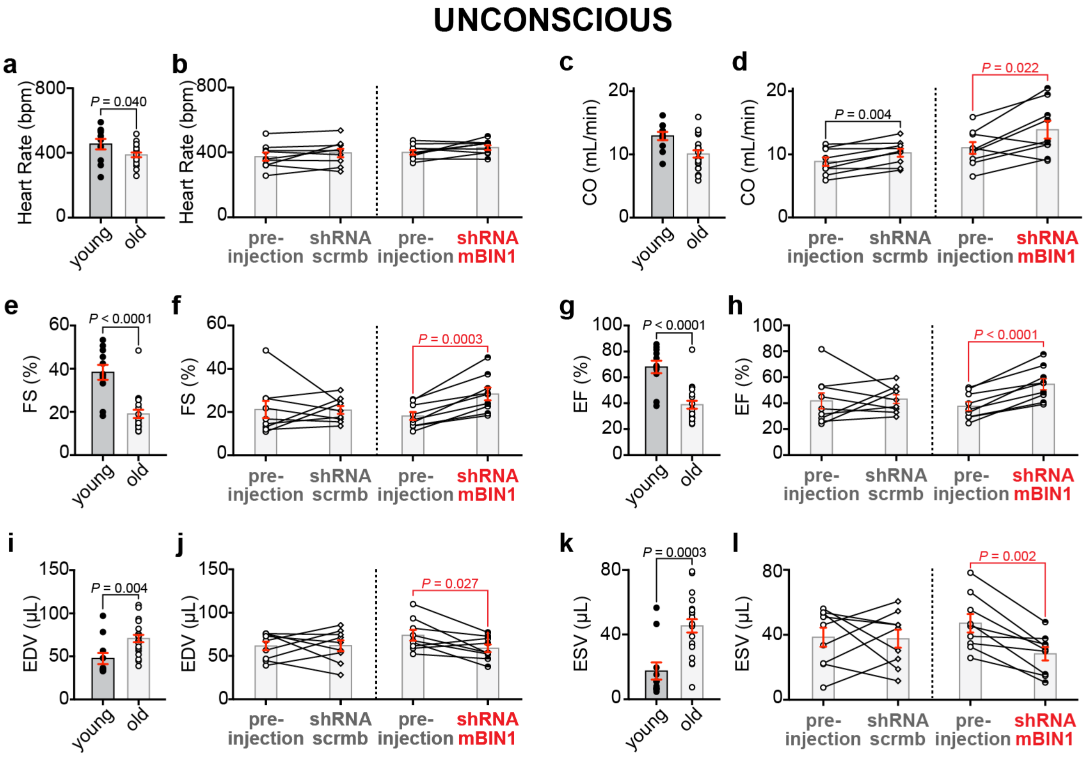


**Supplementary Figure 6. Additional data from unconscious M-mode echocardiography.** Summary dot-plots for unconscious young (*N* = 11) and old (*N* = 20) mice, and paired results before and after RO-injection of old mice with shRNA-scrmb (*N* = 9) and shRNA-mBIN1 (*N* = 9) for the following measurements are displayed: **a** and **b**, heart rate (HR), **c** and **d**, cardiac output (CO), **e** and **f**, fractional shortening (FS), **g** and **h**, ejection fraction (EF), **i** and **j**, end diastolic volume (EDV) and, **k** and **l**, end systolic volume (ESV). Unpaired Student's t-tests were performed on data displayed in a, c, e, g, i, and k. Paired Student's t-tests were performed on data displayed in b, d, f, h, j, and l.

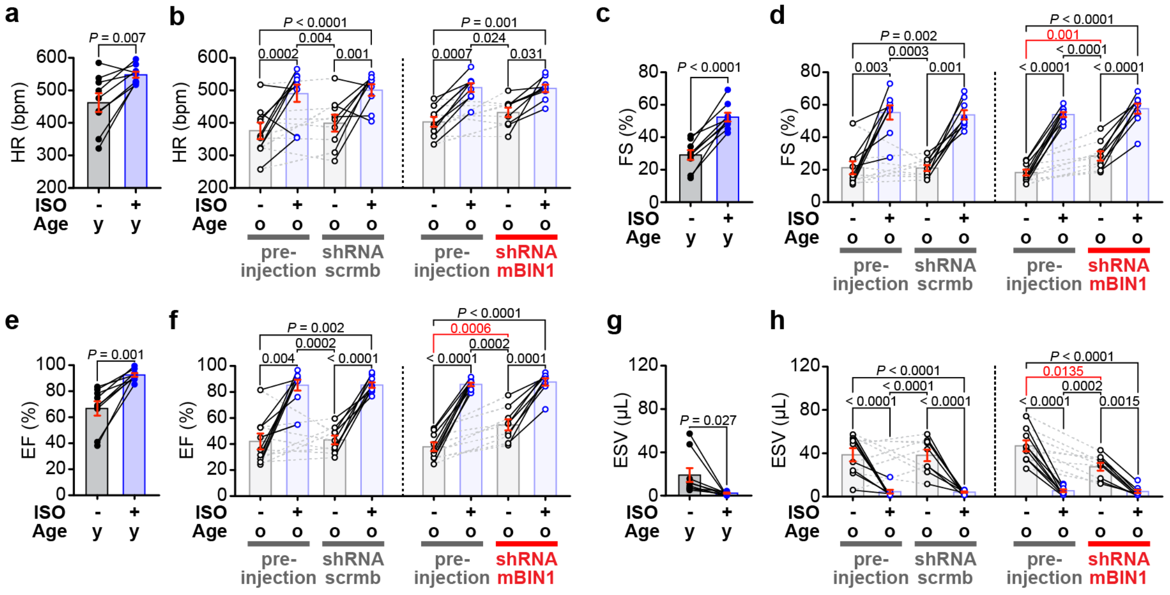


**Supplementary Figure 7. Systolic analysis of *in vivo* ISO response.** Summary dot-plots for unconscious young mice ($N = 9$), and paired results before and after RO-injection of old mice with shRNA-scrmb ($N = 9$) and shRNA-mBIN1 ($N = 9$), before and after intraperitoneal injection of 0.1 mg/kg ISO for the following measurements are displayed: **a** and **b**, heart rate (HR), **c** and **d**, fractional shortening (FS), **e** and **f**, ejection fraction (EF) and, **g** and **h**, end systolic volume (ESV). Paired Student's t-tests were performed on data displayed in a, c, e, and g. Two-way ANOVAs with post-hoc Tukey's multiple comparisons test were performed on data displayed in b, d, f, and h.

| Group | (*N*, *n*) | Fold-change in peak $I_{Ca}$ with ISO | Capacitance (pF) | $V_{1/2}$ (mV) | | Slope factor | |
|---|---|---|---|---|---|---|---|
| | | | | Control | ISO | Control | ISO |
| **WT male young** | *N* = 6, *n* = 8 | 1.59 ± 0.09 | 160.4 ± 13.9 | -11.7 ± 1.0 | -26.5 ± 1.9 *** | 4.8 ± 0.4 | 3.3 ± 0.5 * |
| **WT male old** | *N* = 5, *n* = 11 | 1.19 ± 0.06*** | 153.3 ± 19.9 | -21.1 ± 1.0 | -27.9 ± 1.1 **** | 3.9 ± 0.2 | 3.2 ± 0.2 **** |
| **shRNA-scrmb male old** | *N* = 3, *n* = 7 | 1.14 ± 0.06*** | 162.1 ± 20.4 | -20.8 ± 2.6 | -27.9 ± 2.0*** | 4.9 ± 0.5 | 4.1 ± 0.5* |
| **shRNA-mBIN1 male old** | *N* = 3, *n* = 9 | 1.19 ± 0.06** | 168.7 ± 12.9 | -21.2 ± 1.0 | -30.4 ± 1.0**** | 4.0 ± 0.3 | 3.0 ± 0.3*** |

**Supplementary Table 1. ISO-stimulated changes in peak $I_{Ca}$ and voltage dependence of $G/G_{max}$ of young and old ventricular cardiomyocytes.** Mean ± SEM for the fold-change in peak $I_{Ca}$ (a one-way ANOVA with Tukey's multiple comparisons post-hoc test was used to compare the four groups; * indicates significant difference from young mice), the whole cell capacitance, and for the $V_{1/2}$ and slope factor of $G/G_{max}$ fits (paired Student's t-test; * indicates significant difference between ISO values and their respective controls). This data is also highlighted in Figs. 1 and 5, and Supplementary Fig. 3. *P*-values are represented by asterisks due to space constraints as follows: $^{*}$ $P< 0.05$, $^{**}$ $P < 0.01$, $^{***}$ $P < 0.001$, $^{****}$ $P < 0.0001$.
